# Allele frequency-free inference of close familial relationships from genotypes or low depth sequencing data

**DOI:** 10.1101/260497

**Authors:** Ryan K Waples, Anders Albrechtsen, Ida Moltke

**Affiliations:** Department of Biology, Section for Computational and RNA Biology, University of Copenhagen, 2200 Copenhagen N, Denmark

## Abstract

Knowledge of how individuals are related is important in many areas of research and numerous methods for inferring pairwise relatedness from genetic data have been developed. However, the majority of these methods were not developed for situations where data is limited. Specifically, most methods rely on the availability of population allele frequencies, the relative genomic position of variants, and accurate genotype data. But in studies of non-model organisms or ancient human samples, such data is not always available. Motivated by this, we present a new method for pairwise relatedness inference, which requires neither allele frequency information nor information on genomic position. Furthermore, it can be applied not only to accurate genotype data but also to low-depth sequencing data from which genotypes cannot be accurately called. We evaluate it using data from a range of human populations and show that it can be used to infer close familial relationships with a similar accuracy as a widely used method that relies on population allele frequencies. Additionally, we show that our method is robust to SNP ascertainment and applicable to low-depth sequencing data generated using different strategies, including resequencing and RADseq, which is important for application to a diverse range of populations and species.

## Introduction

The ability to infer the familial relationship between a pair of individuals from genetic data plays a key role in several research fields. In conservation biology, it is used to design breeding programs that minimize inbreeding (Kardos *et al*. 2015), in archeology it is helpful to understand burial patterns and other cultural traditions (Baca *et al*. 2012; Sikora *et al*. 2017), and in population and disease genetics it is often used to exclude relatives, because many analysis methods within those fields assume all analyzed individuals are unrelated and violations of this assumption can lead to wrong conclusions (Balding 2006).

Numerous pairwise relatedness inference methods have been developed, e.g., Thompson (1975), Lee (2003), Purcell *et al*. (2007), Albrechtsen *et al*. (2009), Manichaikul *et al*. (2010), Stevens *et al*. (2011), Korneliussen and Moltke (2015), Conomos *et al*. (2016), Dou *et al*. (2017), and many are available in popular software packages, like PLINK (Purcell *et al*. 2007), and KING (Manichaikul *et al*. 2010). Most of these methods estimate either the three relatedness coefficients *k*_0_, *k*_1_ and *k*_2_, or the kinship coefficient 
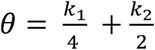
 for each pair of diploid individuals, where *k*_0_, *k*_1_ and *k*_2_ are the proportions of the genome where a pair of individuals share 0, 1, or 2 alleles identical by descent (IBD) (Thompson 2000). By definition, alleles are IBD when they are identical due to *recent* common ancestry, but because alleles can also be identical due to older common ancestry or recurrent mutations, IBD status cannot be directly observed. Therefore, the pairwise relatedness coefficients and the kinship coefficient have to be estimated, which can be done from patterns of observed genetic identity (identity by state; in short IBS). Once estimated, these relatedness statistics, *k*_0_, *k*_1_, *k*_2_, and the kinship coefficient can be used to infer familial relationships by comparison to the expectation of the statistics for different familial relationships (Hill and Weir 2011). Although inference of relatedness is of wide interest, most existing methods are not immediately applicable in studies with limited data or genetic resources. First, most existing methods require the allele frequencies of the source population. For most studies in modern humans this is not a problem. However, in studies of ancient humans or other species, accurate estimates of population allele frequencies are often not obtainable, because only a low number of samples are available. Second, several existing methods consider consecutive loci jointly and use sliding windows or hidden Markov models to leverage the non-independence of allele sharing along the genome between relatives (Albrechtsen *et al*. 2009; Gusev *et al*. 2009; Stevens *et al*. 2011; Kuhn *et al*. 2018). These methods are powerful but require information about the genomic position of variable sites, and for non-model organisms high-quality reference genome assemblies are often not available. Third, other methods avoid allele frequencies, but rely on access to many samples to provide a necessary context for relationship classification (e.g., Abecasis *et al*. 2001). Finally, nearly all existing methods -both frequency-based and others-require genotype data. However, in sequencing studies, samples are often only sequenced to low depth due to cost and technical issues. This makes it infeasible to call genotypes accurately (Nielsen *et al*. 2012), precluding the use of these methods. There are a few methods that estimate relatedness from low-depth sequencing data by utilizing genotype likelihoods (e.g., Korneliussen and Moltke 2015), or by using imputed genotype dosages (Dou *et al*. 2017). However, these methods function by leveraging access to many samples to estimate allele frequencies or perform accurate genotype imputation and are therefore not designed to apply to datasets with a low number of samples.

Hence, most existing methods to infer close familial relationships are not immediately applicable in studies where data is limited, including many studies of non-model organisms and ancient humans. One of the few exceptions is a simple but elegant test for pairwise relatedness proposed in Lee (2003). This test relies entirely on the relative frequency of different genotype combinations within a pair of individuals and thus only requires genotype data from the two target individuals. While useful, this test does not provide any means to distinguish between different types of close familial relationships; it only provides a statistical test for a pair of individuals of the null hypothesis of them being unrelated.

There are only a few methods that can be used to distinguish between different types of familial relationships for a pair of individuals when neither allele frequencies nor information about the relative genomic position of sites is obtainable. One such method consists of plotting the proportion of the genomic sites in which the two individuals share both alleles IBS (which we will denote IBS2) versus the proportion of the genomic sites in which they share zero alleles IBS (which we will denote IBS0). This method was used in Rosenberg (2006), where it was applied to the Human Genome Diversity Project (HGDP) dataset. In the resulting scatter plot of the HGDP data (figure 1 of Rosenberg (2006)), pairs of individuals with the same relationship category form distinct clusters so that it is possible to locate parent-offspring pairs, full sibling pairs, and to a lesser extent more distant relationships such as half-siblings/avuncular/grandparent-grandchild and first cousins.

**Figure 1.**
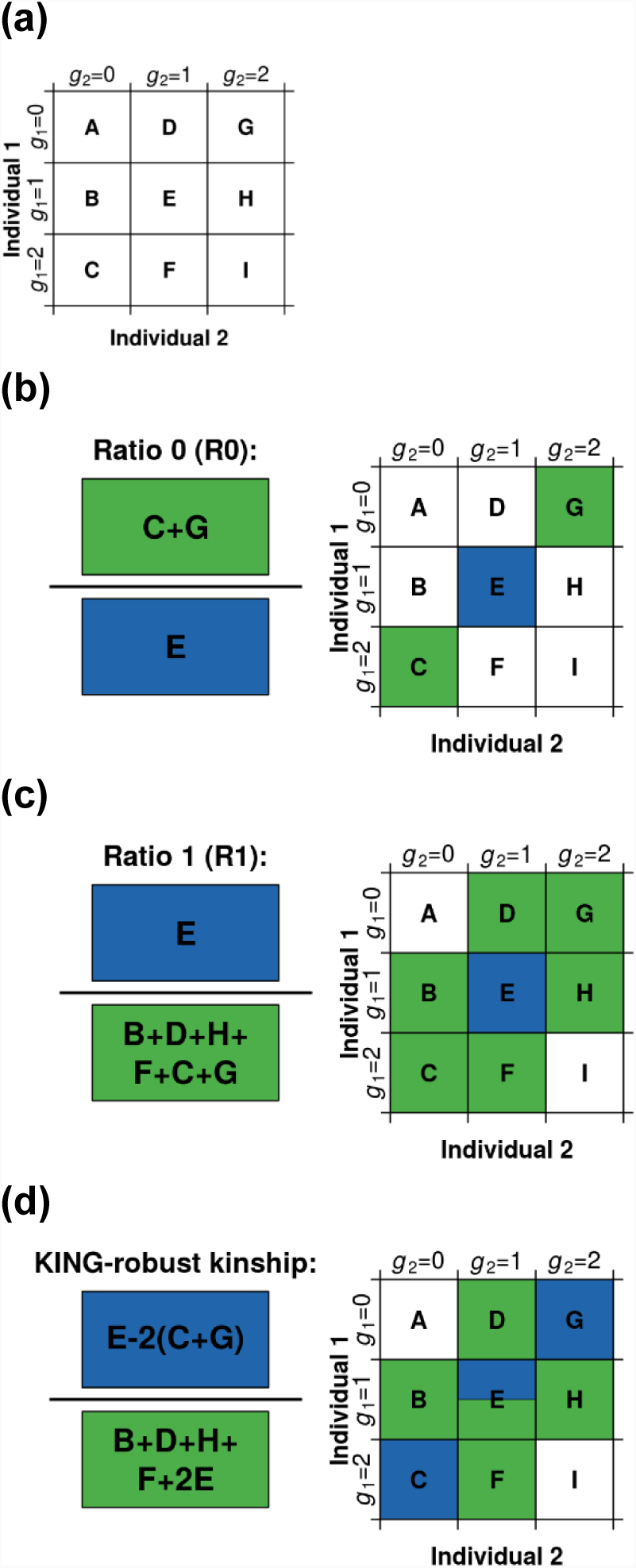
Definitions of pairwise genotype categories A - I and the R0, R1, and KING-robust kinship statistics. **(a)** Definition of the pairwise genotype categories A - I. Here g_1_ and g_2_ denotes the genotype for each of the two diploid individuals, 1 and 2, respectively. These genotypes are defined as the number of copies of a certain allele carried by 1 and 2, respectively. We assume diallelic variants such that g_1_ and g_2_ each has 3 possible values: 0, 1, and 2. For a pair of individuals, there are nine possible genotype combinations. We organize them into a 3x3 matrix and denote them with the letters from A to I. The values A – I can equivalently be either counts or proportions. **(b)** Definition of the R0 statistic based on the notation illustrated in A. **(c)** Definition of the R1 statistic based on the notation illustrated in A. **(d)** Definition of the KING-robust kinship estimator (Manichaikul *et al*. 2010), formulated using the notation illustrated in (a).

Another such method is based in part on the KING-robust kinship estimator (Manichaikul *et al*. 2010). The KING-robust kinship estimator was developed to be robust to population structure, but in practice it has been shown to provide biased kinship estimates when applied to pairs of samples whose four chromosomes are not all from the same population (Thornton *et al*. 2012; Conomos *et al*. 2016). However, the KING-robust kinship estimator is directly applicable to samples from the same homogenous population even when allele frequencies are unknown. The reason for this is that, like the test suggested in Lee (2003), it relies only on the genotype combinations within the two target individuals and does not require knowledge about of allele frequencies. Importantly, Manichaikul *et al*. (2010) show it is possible to infer if a pair of individuals are parent-offspring, full sibling, half siblings/avuncular/grandparent-grandchild, first cousins, or unrelated, by jointly considering KING-robust kinship and the fraction of sites IBS0 using SNP array data without allele frequencies. For example, see figure 3A in Manichaikul *et al*. (2010); a scatterplot of KING-robust kinship vs the fraction of sites IBS0.

However, both the methods described above have two important limitations. First, like most other methods to estimate relatedness, they were developed for genotype data only. For example, the KING software (Manichaikul *et al*. 2010) implementing the KING-robust kinship estimator requires genotype data as input, which can be problematic for studies where only moderate or low-depth sequencing data is available and calling genotypes is consequently difficult. Second, both methods rely on estimates of the fraction of sites IBS0, which can be problematic because this fraction, as well as the fraction of sites IBS1 and IBS2, is highly sensitive to SNP ascertainment. This means that the results of the methods are platform-dependent and are likely to differ between different SNP arrays and especially between SNP array and sequencing datasets. And in turn, this among other things, means that it can be difficult to distinguish between full siblings and parent-offspring pairs using these methods.

Motivated by the outlined limitations to the existing methods, we present a method for relationship inference that -unlike most existing methods-relies neither on allele frequencies nor on information about the relative position of the variant sites, and which – unlike other frequency-free methods – 1) is applicable even to sequencing data of so low depth that accurate genotypes cannot be called from it and 2) is robust to SNP ascertainment bias.

The new method is inspired by previous methods; it uses the KING-robust kinship estimator and a statistic R0, which is similar to the test statistic from the test for relatedness suggested by Lee (2003). However, the method is new in two important ways. First, besides relying on the two statistics, R0 and KING-robust kinship, it also relies on a third new statistic, R1. More specifically, the method consists of using two combinations of these three statistics, R1-R0 and R1-KING-robust kinship, to infer relationships, and it is this combination of statistics that makes the method robust to ascertainment bias. Secondly, while the new method can straightforwardly be applied to genotype data like other similar methods, we also present two computational approaches to estimate the three statistics directly from sequencing data that takes the uncertainty of genotypes into account, allowing application to low-depth sequencing data.

In the following we first fully describe the three statistics, R0, R1, and KING-robust kinship, how they can be estimated and other methodological details. Next, using simulated and publicly available SNP array data, we show that the new method provides similar accuracy and precision to the commonly used frequency-based method implemented in PLINK, when such data is available. Then using sequence data from the 1000 Genomes Project (The Genomes Project 2015), we show that the three statistics can be estimated directly from sequencing data of low depth (∼4x), here defined as depth insufficient for accurate genotype calling. Moreover, we show that the estimates obtained in this way are useful for inference of close familial relationships and that this is not the case for estimates obtained from genotypes called from the same data. Using different subsets of the same data, we also show that this new method, unlike previous similar methods, is robust to SNP ascertainment. Finally, we show that the method also provides useful results when applied to sequencing data down-sampled to approximate data generated using reduced-representation approaches, e.g. restriction-site associated DNA sequencing (RADseq) and discuss some potential applications and limitations of the new method.

## Methods and materials

### The R0, R1, and KING-kinship statistics

The method for relationship inference we propose consists of estimating three statistics called R0, R1 and KING-robust kinship from genetic data and interpreting plots of R1 vs R0 and R1 vs KING-robust kinship.

We define the three statistics, R0, R1, and KING-robust kinship in terms of the genome-wide IBS sharing pattern of two individuals. At any given diallelic sites, a pair of individuals will carry one of nine possible genotype combinations; the nine possible combinations of the two individuals carrying 0, 1, or 2 copies of a specific allele, e.g., the ancestral allele. We can therefore fully characterize the genome-wide IBS-sharing pattern of a pair of individuals by nine counts or proportions denoted: A, B, C, D, E, F, G, H, and I (Figure 1A), similar to a two-dimensional site-frequency spectrum (SFS) across the two individuals. The R0 and R1 statistics are defined as simple functions of a subset of these nine counts as shown in Figure 1B and C, and the KING-robust kinship statistic, originally defined by Manichaikul *et al*. (2010), can also be re-formulated as a function of these 9 values (Figure 1D).

The new method is motivated by several observations. First, the expected values of A-I vary depending on the familial relationship between the pair of individuals of interest. Consequently, so do functions of A-I, including R0, R1, and KING-robust kinship. Notably, there is no overlap between the joint expectation ranges of [R1, R0] and [R1, KING-robust kinship] for the four close relationship categories: full siblings (FS), half siblings/avuncular/grandparent-grandchild (HS), first cousins (C1) and unrelated (UR) and the range of expected values for parent-offspring (PO) only overlaps with those of FS in a single point (Figure 2, for derivations see supplementary text). Crucially, this is true regardless of the underlying allele frequency spectrum and holds for any pair of non-inbred individuals from the same homogenous population, making [R1, R0] and [R1, KING-robust kinship] potentially useful for distinguishing between these relationships. Second, while A-I, and thus R0, R1, and the KING-robust kinship estimator can be calculated from genotype data, they can also be estimated directly from next-generation sequencing (NGS) data based on the expected number of sites with each genotype combination (see below for details). This makes the method appropriate even when the sequencing depth is too low for accurate genotype calling (see below for methodological details). Third, regardless of the type of data that is available, R0, R1 and KING-robust kinship can be estimated without the need for population allele frequencies and information about the relative position of the genomics sites analyzed. Finally, we expect the three statistics to be robust to SNP ascertainment because they are ratios computed from sites that are variable within the two samples and should thus be unaffected by the number of non-variable sites and because the (unknown) underlying frequency spectrum should only have a limited effect on these ratios.

### Estimation from sequencing data

The counts of the nine genotype combinations, A-I, and thus R0, R1, and KING-robust kinship for a pair of individual, can be estimated directly from NGS data via the use of genotype likelihoods calculated from the aligned sequencing reads. Genotype likelihoods provide a means to account for the genotype uncertainly inherent to low-depth NGS data. We used two distinct, but related, approaches to estimate these statistics from sequencing data that both build on this idea.

The first approach, which we denote the IBS-based approach, considers all ten possible genotypes at a diallelic site for each of the two individuals of interest and consists of a maximum-likelihood (ML) estimation of the counts of each of the 100 (10 × 10) possible genotype pairs (for details see supplementary text). To perform the ML estimation, we used an EM algorithm, which we have added to the ANGSD software package under the name IBS. After obtaining the estimate of the counts of all the 100 possible genotype pairs, we convert them into estimates of A-I, by summing over the counts that correspond to each combination. For example, the genotype pairs: AA/AA, CC/CC, GG/GG and TT/TT all contribute to cells A or I of Figure 1. Counts corresponding to genotype pairs with more than two different alleles (e.g. AC/AG) were discarded. The advantage of this IBS-based approach is that it does not require specification of a known allele at each site and can thus be applied to nearly any sequencing data set, even with low-depth, without any prior information on the alleles at each site.

The second approach, which we denote the SFS-based approach, consists of performing ML estimation of the two-dimensional site-frequency spectrum (SFS) (Nielsen *et al*. 2012). To find the ML estimate of the SFS, we used an expectation-maximization (EM) method implemented in the ANGSD software package under the name realSFS (Korneliussen *et al*. 2014). The SFS-based approach requires one allele to be specified for each site e.g., the ancestral, consensus, or reference allele. The model underlying this approach assumes that genotypes for each site have the possibility of containing this specified allele and up to one other unspecified allele. To specify the alleles that exist at each site, we used the consensus sequences from the highest depth individual (NA19042) and restricted our analysis to sites where the depth in this individual was at least three.

The computational burden of analyzing genome-wide data sets can be significant. For both the genotype likelihood-based approaches described above, the main limitation is RAM, as data likelihoods for each site need to be loaded into memory for optimization. To overcome this limitation, we analyzed each chromosome separately and then summed over the values for each chromosome to produce a genome-wide estimate. To calculate the likelihood of sequencing data for each possible genotype, we used the original GATK genotype likelihood model (Mckenna *et al*. 2010) with independent errors, as implemented in ANGSD. We also tried to use the samtools genotype likelihood model (Li 2011) with a more complicated error structure, but found it produced worse results (data not shown). Both IBS and realSFS produce the expected values for A-I, which we subsequently use to calculate R0, R1 and KING-robust kinship. For example command lines for each analysis, see supplemental text.

### Confidence intervals

All of the above estimation methods treat each site as independent. This assumption should not affect our expectation of each statistic (Wiuf 2006), but statistical non-independence (here due to linkage disequilibrium (LD) and IBD) does affect standard estimates of uncertainty. To quantify the uncertainty, we therefore estimated confidence intervals for all statistics using a block-jackknife procedure. Confidence intervals were estimated by leaving each chromosome out (chromosome jackknife), which takes both the IBD and the LD correlation into account. The weighted block jackknife variance estimator of Busing *et al*. (1999) was used to estimate the variance from the distribution of estimates for each statistic. The square root of this variance was interpreted as the standard error in our estimate.

### Application to simulated data

To evaluate the new method and to investigate the effects demography, we simulated genotype data under three different demographic histories: 1) a neutral demography with constant effective population size (*N_e_*), 2) a shrinking demography with a 10x decrease in *N_e_* over the last 100 generations, 3) an expanding demography with a 10x increase in *N_e_* over the last 100 generations. For each of the three scenarios we used the coalescent simulator `msprime` (Kelleher *et al*. 2016) to simulate four haplotypes. These haplotypes were then used to construct pairs of related individuals in five relationship categories: parent offspring (PO), full siblings (FS), half-siblings/avuncular/ grandparent-grandchild (HS), first cousin (C1), and unrelated (UR). For unrelated pairs of individuals, the genotype data were constructed by simply splitting the four haplotypes into two pairs. For related pairs of individuals, we constructed the genotypes at each variable site by first sampling whether the individuals shared 0, 1, or 2 alleles IBD according to the expected values of [k0, k1, k2] for the relevant relationship (Supplementary Table S1). Then we used alleles present on the four haplotypes according to the sampled IBD status to construct the genotype at that site. For example, if the individuals were selected to share two alleles IBD at a site, both individuals received the alleles present on the first two haplotypes. The IBD sharing pattern was sampled independently for each SNP and thus LD and biological variation in IBD was not modelled. We concatenated the data from many independent simulations to achieve enough data so the IBD sharing was approximately equal to the expected values. Simulation code is available in the supplemental materials.

### Application to real datasets

To assess the utility of the new method on more realistic data, we applied it to two different publicly available datasets: SNP array data from seven HGDP populations (Rosenberg 2006), and sequencing data from five related individuals from the Luhya in Webuye, Kenya (LWK) population of the 1000 Genomes Project phase 3 (The Genomes Project 2015).

### HGDP SNP array

The HGDP SNP array dataset was accessed Jan. 13, 2017 and we followed the quality control steps described in Rosenberg (2006) to exclude mislabeled and duplicate samples. We selected seven populations from the HGDP based on the presence of several close familial relationships; five non-African populations: Surui, Pima, Karitiana, Maya, and Melanesian, and two African populations: Mbuti Pygmies and Biaka Pygmies.

To ensure a fair comparison to the allele frequency based inference method in PLINK, and by proxy to other commonly used methods, we constructed data sets where these methods have been shown to perform well. Specifically, we excluded individuals showing obvious signs of admixture (n=16) or inbreeding (n=2) from the selected HGDP populations. For details see Supplemental text 2.3.1. This left us with a total of 142 individuals from the seven populations: Surui N=20, Pima = 20, Karitiana N=21, Maya N=16, Melanesian N=19, Biaka Pygmies N=31 and Mbuti Pygmies N=15. For each of these seven populations we constructed a final set of genotypes by retaining genotypes from autosomal loci with genotyping rate >99%, minor allele frequency (MAF) >5%, HWE p-value > 10^-4^.

The Ro, R1, and KING-robust kinship statistics for each of the 2902 within-population pairs of individuals were then calculated from all sites where both individuals had non-missing genotypes.

### 1000 Genomes sequencing data

To get sequencing data from several different relationship categories, we selected five individuals from two families in the Phase 3 1000 Genomes (1000G) Luhya in Webuye, Kenya (LWK) population: NA19027, NA19042, NA19313, NA19331, NA19334. Across the five individuals, there is one pair of half-sibs (NA19027 & NA19042), and a separate trio of related individuals with a pair of full siblings (NA19331 & NA19334), one parent-offspring relationship (NA19313 & NA19331), and another unspecified second-degree relationship (NA19313 & NA19334), possibly avuncular (The Genomes Project 2015). These stated relationships leave six unrelated pairs among the five individuals.

For each pair of the five LWK individuals we estimated the R0, R1, and KING-robust kinship statistics in five different ways: 1&2) by applying the two different sequencing-based approaches described above to the 1000G aligned sequence data files (∼4x coverage bam files), 3) by simple genotype counting based on the phased and high-quality curated genotypes provided in the hg37 1000G VCF files, 4) genotype counting based on the subset of sites in 3 that overlap with the Illumina 650Y sites for the HGDP data (to investigate ascertainment, see below) and 5) by calling genotypes from the same 1000G bam files in a basic manner meant to mimic data from a species with a reference genome but few other genetic resources and then simply counting from the called genotypes. For set 5, we used samtools mpileup (v1.3.1) to summarize the reads overlapping each position, and bcftools call (v1.3.1) to assign the most likely genotype at each position. We used mostly default settings; non-default flags to samtools specified skipping indel positions. Non-default flags to bcftools specified using the consensus caller. For all sequence-based analyses (1,2 &5) we only considered reads with a minimum phred-scaled quality score of 30 and bases with minimum phred-scaled quality score of 20 and we restricted our analyses to genomic regions with a GEM 75mer mappability of 1 (Derrien *et al*. 2012). Notably, all the methods and filters used here can be applied to any study, including studies with only small contigs, for example made up of RAD loci, making the results relevant beyond resequencing studies utilizing well assembled genomes.

### Assessing the effect of SNP ascertainment

To evaluate the effect of SNP ascertainment using real data, we created a subset of the curated genotype data from the five 1000 Genomes individuals. We selected the sites that overlap with the Illumina 650Y array that was used for the HGDP and estimated our three relatedness statistics. We compared the results from this subset of HGDP sites to results for the full genotype dataset and also to the sequence-based analyses. For an additional comparison, we also performed the same comparison for the methods presented in Rosenberg (2006) and Manichaikul *et al*. (2010) by constructing scatterplots by their methods.

We also investigated the effect of SNP ascertainment using the data simulated from a constant demography (see “Application to simulated data” for details about the simulations) and compared results for the full dataset to results obtained by including only sites with a minor allele count >2 out of 40 chromosomes (MAF >5%) in the analyses.

### Assessing the effect of a limited number of sites

To assess the usefulness of this new method on data sets with fewer genomic sites covered by sequencing reads we constructed reduced size data sets, in a way that mimicked some aspects of reduced-representation sequencing approaches such as RADseq. To produce each reduced data set, we selected a specific number of 200bp windows randomly from the eligible genomic regions and restricted our analysis to sites falling within them. We used 10k 50k, 100K, and 250k windows, representing ∼4x sequencing coverage on 2M, 10M, 20M, or 50M sites, respectively. All other aspects of the analyses were the same, expect for that for these data sets, we applied the IBS- and SFS-approaches to the complete data set, rather than splitting by chromosome as we did for the full dataset. We suggest analyzing the complete data in single run if your computational resources allow it, as we noticed some upward bias in the estimated number of IBS0 sites when the smaller data sets were analyzed separately by chromosome.

### Comparison to other methods

To get a categorization of relationships for the HGDP dataset described above based on a standard, commonly used allele frequency-based method, we first applied the allele frequency based relatedness estimation program PLINK (v1.9) (Chang *et al*. 2015) to the individuals from each population separately to estimate the genome-wide IBD fractions *k*_0_, *k*_1_, *k*_2_. Next, we applied the relationship criteria proposed in Table 1 of Manichaikul *et al*. (2010) to the relatedness coefficient estimates: the estimated k values were combined into an estimate of the kinship coefficient 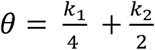, and a relationship degree was assigned to each pair of individuals based on comparing the estimated kinship coefficient to the criteria in the table. Parent-offspring and full siblings were differentiated based on *k*_2_ values. This provided us with a categorization into five categories: PO, FS, HS, C1, and UR. To achieve additional resolution, we further divided the last category (UR) into two: unknown/distantly related (UK-DR) and unrelated (UN). We did this by simply extending the logic behind the criteria proposed above. Specifically, we set the kinship threshold between UK-DR and UR to 1/2^13/2^, which corresponds to including 4^th^ to 5^th^ degree relatives in the UK-DR category.

To assess the accuracy and precision of the new method for familial relationship classification within the HGDP data we examined concordance with the PLINK-based relationships described above. For this purpose, we assigned a relationship category to each pair of individuals in two ways: 1) using the statistics R0 and R1 and 2) using a combination of KING-robust kinship and R0. For the former, we characterized each possible relationship by a single [R1, R0] point generated from data simulated under a demography with a constant population size over time, detailed in the “Application to simulated data” section and assigned each pair of individuals the relationship of the closest point using a Euclidean distance measure. For the latter, we first used the KING-robust kinship criteria from table 1 of Manichaikul *et al*. (2010) as above. Since this table has overlapping kinship ranges for the PO and FS categories, we used the R0 statistic to distinguish PO from FS relationships: Ignoring rare effects like germline mutations and genotyping errors the expected value for R0 for PO relatives is zero, while for FS the value is above 0; we used an ad hoc cut-off of 0.02.

To estimate the statistics for identifying related individuals proposed by Rosenberg (Rosenberg 2006) and KING (Manichaikul *et al*. 2010), we note that the KING-robust kinship estimator can (as previously described) be calculated directly from the same nine counts, A-I, and so can the fraction of sites IBS0 and IBS2:

KING-robust kinship = (E – 2(C+G))/ (B+D+H+F+2E)

Fraction IBS0 = (C+G)/(A+B+C+D+E+F+G+H+I)

Fraction IBS2 = (A+E+I)/(A+B+C+D+E+F+G+H+I)

We used these formulas in all our comparisons because this allowed us to estimate these statistics not only from genotype data but also directly from sequence data in the same manner as for R0 and R1. However, we note that this is our approach to estimating those statistics, and that existing tools like KING only allow users to estimate the statistics from genotype data.

## Results

To assess the performance of the new method, we first applied it to simulated genotype data to ensure that it works on sufficient data and to assess how sensitive it is to the underlying demographic history of the population the analyzed samples are from. Next, we applied the new method to real data from different platforms to assess its performance on more realistic data. Finally, we performed a couple of additional analyses to access how robust the new method is to SNP ascertainment and to having data from only a limited number of sites available. Below we describe the results of all these analyses.

### Application to simulated data

We first applied the new method to simulated genotype data from several different relationship pairs from populations with three different demographic histories: 1) constant *N*_*e*_, 2) 10-fold increase in *N*_*e*_ over the past 100 generations, and 3) 10-fold decrease in *N*_*e*_ over the past 100 generations. We see very similar results across all three demographic scenarios and in all cases the R1-R0 and R1-KING-robust kinship values obtained from the simulated data were within the theoretically derived ranges of expectations (Figure 2). These results demonstrate that the method works if sufficient high-quality genotype data is available. Furthermore, they demonstrate that the range of examined population size histories has a limited effect, even though demographic history affects the allele frequency spectrum. In turn, this suggests that the range of expected values that are realistic for real data is markedly smaller than the theoretically derived ranges also shown in the figure, which is useful for classification purposes.

**Figure 2.**
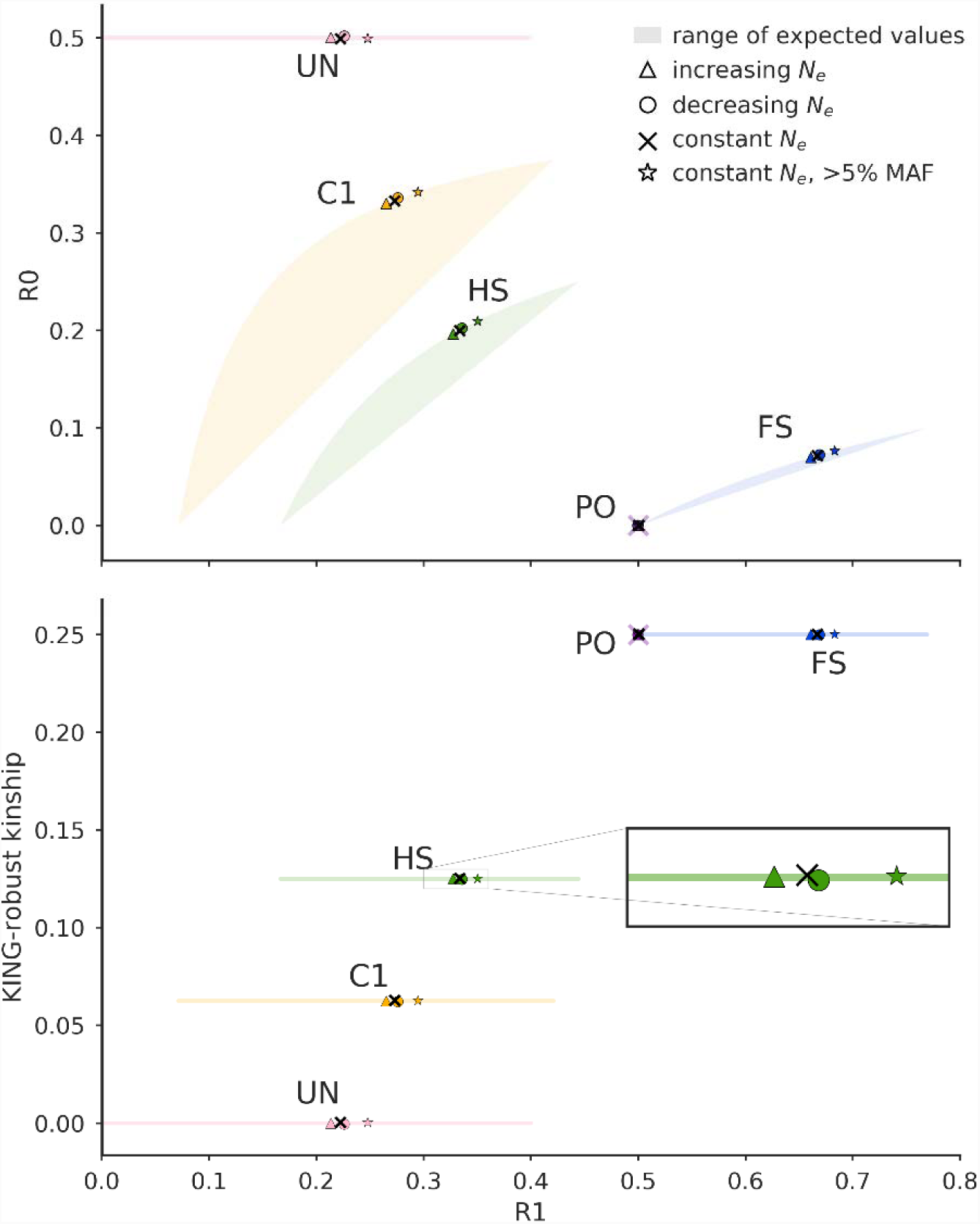
Ranges of expected values and simulation results for R1-R0 and for R1-KING robust kinship for each of five relationship categories: parent-offspring (PO), full siblings (FS), half siblings/avuncular/grandparent-grandchild (HS), first cousins (C1), and unrelated (UR). **Top)** The colored shaded areas (sometimes just lines) show theoretically derived ranges of the joint expectation for R1-R0 based on expected IBD sharing (i.e. values of k0, k1 and k2) for each relationship across all possible allele frequency spectra. For PO this range is a singular point and is shown as a shaded purple X. The colored symbols (triangle, circle, star) and black ‘x’s show values for each relationship obtained from data simulated under four different scenarios. Three of the scenarios are different demographic histories: 1) a 10-fold increase in *N_e_* over 100 generations, 2) constant *N*_*e*_, and 3) a 10-fold decrease in *N*_*e*_ over 100 generations. The fourth scenario is also constant *N*_*e*_, but sites are ascertained to have allele frequency above 5%. Note that, while it is difficult to see due to overplotting, all simulated values for PO fall very close to (R1, R0) = (0.5, 0). **Bottom)** same top, but for (R1, KING robust kinship). Here all simulated values for PO fall very close to (R1, KING robust kinship) = (0.5,0.25).

### Application to SNP array data

Next, we applied the method to SNP array data from the Human Genome Diversity Project (HGDP) to see how the method works on real data, for which standard allele frequency-based methods like PLINK are known to perform well. More specifically, we applied it to genotype data from unadmixed and non-inbred samples from seven populations originating from the HGDP. This resulted in the R1-R0 and R1-KING-robust kinship plots shown in Figure 3A and B (for population-specific plots see Supplementary Figures S1 and S2). The true relationships for the pairs of individuals are not known; we instead colored each point in Figure 3A and B according to the relationship category inferred based on results from the standard, commonly used allele frequency-based method PLINK (Figure 3C, Supplementary Figure S3). Since there are least 15 individuals in each of the seven selected HGDP populations, the allele frequency estimates for these populations should be reasonably accurate even with some relatedness. Hence, with the large amount of data available in this dataset, the allele frequency-based method should provide correct inference of most, if not all, pairs closer than first cousins, but may not be able to fully distinguish first cousins from more distantly related pairs.

**Figure 3.**
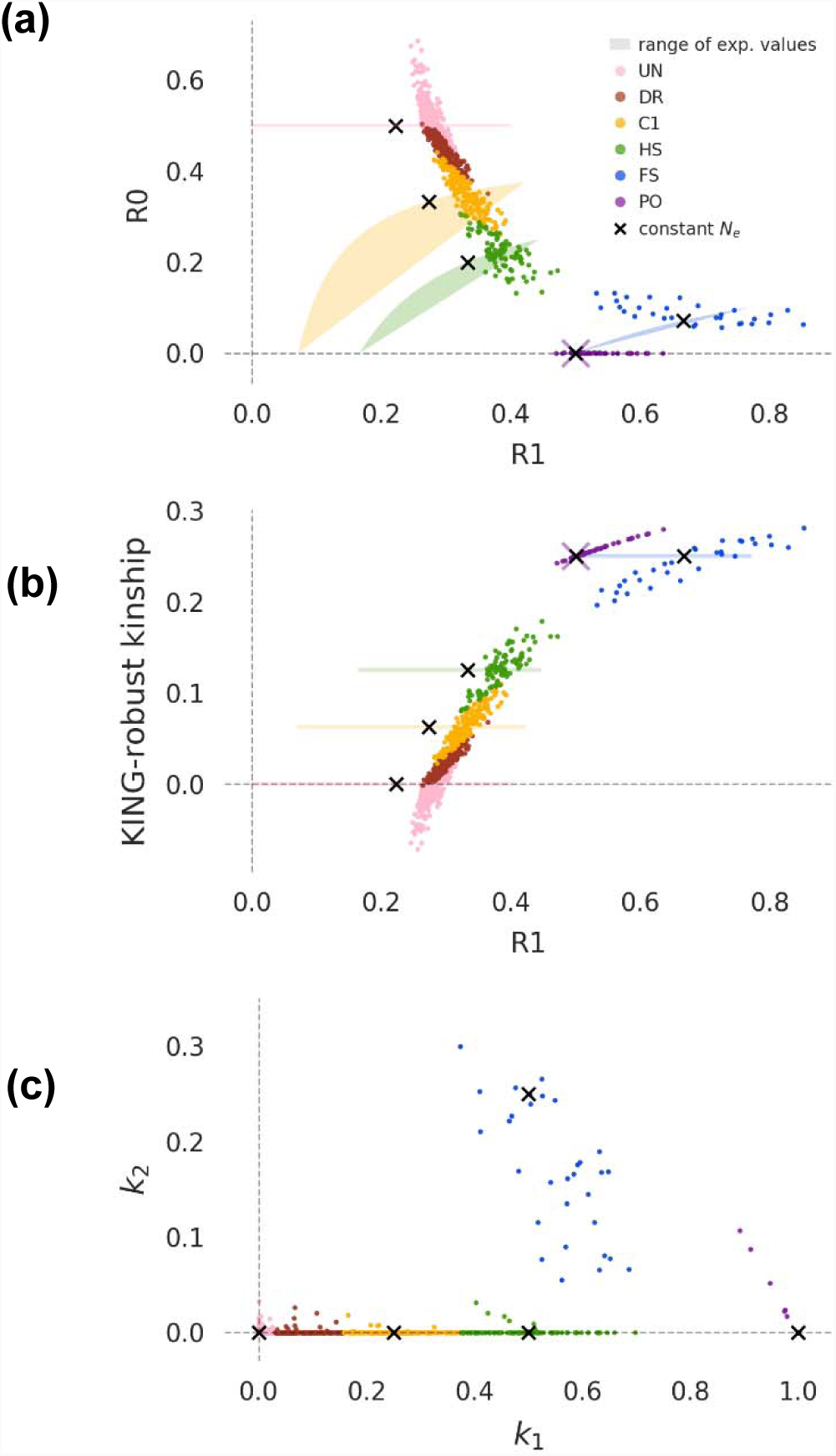
R1-R0 and R1-KING-robust kinship scatterplots for seven HGDP populations. Each colored point represents a pair of individuals and is colored according to the relationship category inferred using an allele frequency based approach. Colored shaded areas/lines show the theoretically derived range of expected values for specific relationship categories as in Figure 2. Black ‘X’s show the values for a pair of individuals simulated under a constant population size demography from Figure 2. Note that in addition to the relationship categories for Figure 2 there is an additional category here representing distantly related pairs (DR). **(a)** R1-R0 plot for all pairs of individuals within each population **(b)** R1-KING-robust kinship plot for all pairs of individuals within each population. **(c)** Scatterplot of the two relatedness coefficients k1 and k2 for all pairs of individuals within each population estimated using the allele frequency-based approach implemented in PLINK. Note that the black ‘X’s here show simulated values for k1 and k2 and are not inferred by PLINK, they approximately coincide with the expected values of k1 and k2 for each relationship category (Supplementary Table S1).

In Figures 3A and B, points from each relationship category clearly cluster together on both the R1-R0 plot and the R1-KING-robust kinship plot. Moreover, these clusters are located near both their theoretically derived ranges of expected values and the values from simulated data. In a similar manner, the *k*_1_-*k*_2_ values for the same pairs of individuals cluster close to the expected and simulated values of *k*_1_ and *k*_2_ (Figure 3C). Almost all pairs identified as parent-offspring (PO) by the frequency-based method are easy to identify as such in both the R1-R0 plot and the R1-KING-robust kinship plot, which is not the case when only a single statistic is used (see also Supplemental Figures S1 and S2). The same is true for full siblings (FS). Furthermore, points classified as half-siblings/avuncular/grandparent-grandchild (HS) or first cousins (C1) by the frequency-based method have a minimal overlap with each other and with less-related pairs (Figure 3A and B). The few pairs of individuals that were difficult to classify are the same pairs as those that are edge cases for the allele frequency-based method. This is apparent in an R1-R0 plot of the HGDP data constructed excluding pairs that are closer than 0.01 to the kinship coefficient thresholds that the frequency-based method used when classifying relationships (Supplementary Figure S4).

To quantify precision and accuracy, we examined the concordance between classifications based on the new method and the PLINK-based classification. We tried two simple classification schemes: one based on R1-R0, which uses proximity to the values we obtained from simulated data from a constant *N*_*e*_ demography, and one based on KING-robust kinship (for details see Methods and Materials). The results supported the visual assessment: both classification schemes are highly concordant with the classifications obtained using the frequency-based method (Supplemental Figure S5). Mean precision across all relationship categories was 0.90 for the R1-R0 method, vs 0.89 for KING-robust kinship. Mean recall across all relationship categories was 0.88 for R1-R0, vs 0.89 for KING-robust kinship. The relationship categories for which the method has the lowest precision is the first cousins versus less related pairs, where the allele frequency-based method is also known to have a hard time making classifications. For PO, FS and UR alone the mean precision is as high as 0.99 for R1-R0 and 0.96 for KING-robust-kinship, and the mean recall for these three categories is as high as 0.96 for R1-R0 and 0.99 for KING-robust kinship. Hence the new method provides comparable performance to a frequency-based method when sufficient genotype data is available, but without the need for allele frequency information.

### Application to sequencing data

To assess how well the new method works on more limited real data, we applied it to sequencing data from five low-depth (∼4x) human genomes from the 1000 Genomes project. Among the five selected samples there is a parent-offspring pair, a pair of full siblings, a pair of half siblings, an unspecified 2^nd^ degree relationship (e.g. avuncular), and the rest are unrelated. We estimated the R0, R1 and KING-robust kinship for each pair in several ways. First, by using an IBS-based approach that estimates the proportion all pairwise combinations of the 10 possible genotypes (Figure 4, “IBS”). Second, by using an SFS-based approach where we estimated the two-dimensional site frequency spectrum (2D-SFS) of each pair with a bi-allelic model and calculated R0, KING-robust kinship, and R1 based on this spectrum (Figure 4, “realSFS”). Both these approaches base their estimates on genotype likelihoods calculated from the sequencing read data, instead of on called genotypes, and take the uncertainty of the underlying genotypes that is inherent to low-depth sequencing data into account. The key difference between them is that the SFS-based approach requires specification of an allele known to exist at each site, whereas the IBS-based approach has no such requirement, making it more generally applicable. The approaches also differ in how they deal with sites with more than two unique alleles, either excluding them (IBS-based approach) or integrating over the two-allele possibilities (SFS-based approach), but these sites are rare (mean fraction as estimated by IBS: 1.8E-6) so the impact of discarding them is minimal.

**Figure 4.**
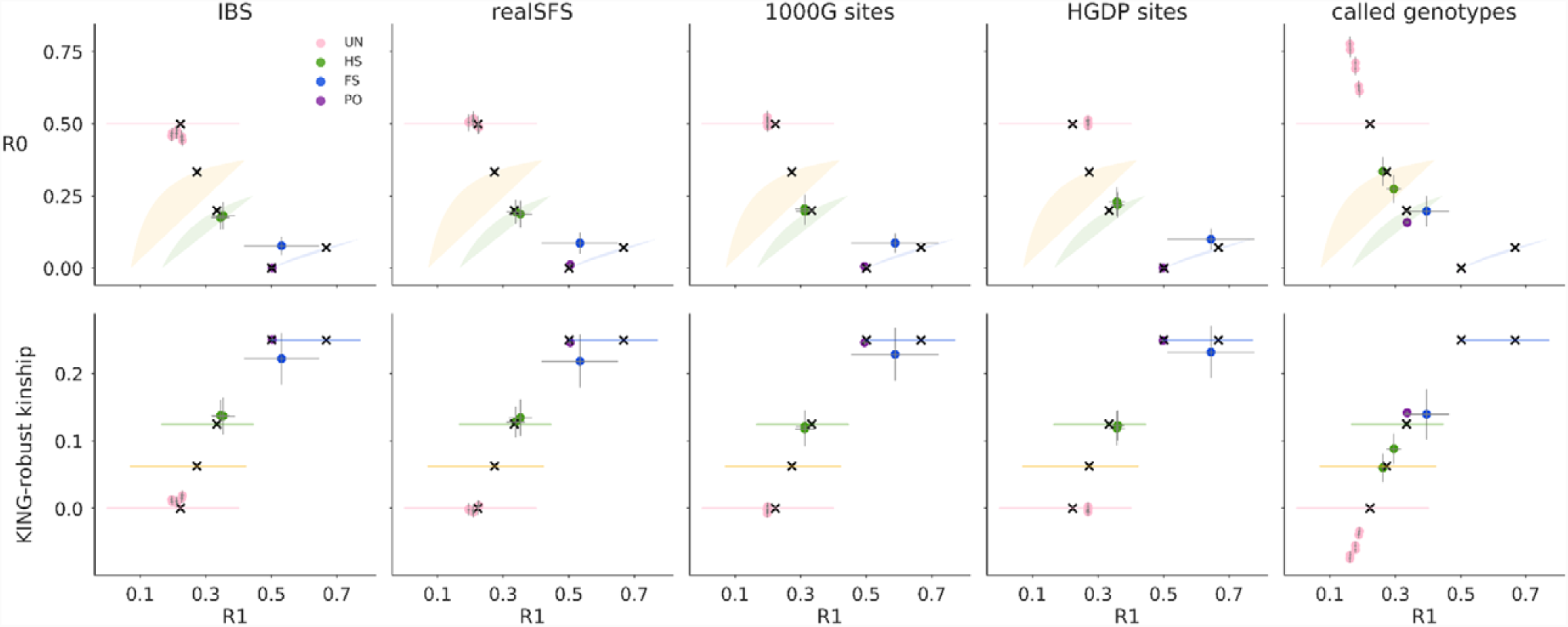
Relatedness plots for all pairs among five LWK individuals from the 1000G Project. **Top)** R1-R0 scatterplots for pairs of five LWK individuals for five different analysis methods: 1) IBS: reference-free estimation from (∼4x) 1000G bam files, 2) realSFS: site-frequency spectrum based estimation from (∼4x) 1000G bam files, 3) 1000G sites: curated 1000G genotypes from the 1000G project, 4) HGDP sites: curated 1000G genotypes but only at sites that overlap with the Illumina 650Y array used for the HGDP, and 5) called genotypes: genotypes called de novo from (∼4x) 1000G bam files‥ Points are colored by their true relationship status, as reported by 1000G. Thin grey lines show confidence intervals (+/-2 SE) estimated using a chromosome jackknife. Colored shaded areas/lines show the theoretically derived range of expected values for specific relationship categories from Figure 2. Black ‘X’s show the values for a pairs of related individuals simulated under a neutral demography, as in Figure 2. **Bottom)** R1-KING-robust kinship scatter plots for the same data sets, confidence intervals and expected ranges are constructed in the same way.

With the values estimated using the SFS-based approach it is possible to visually classify of all the pairs to their relationship category within the set of close familial relationships [PO, FS, HS, or UR], or by using one of the classification methods introduced earlier). Results for the IBS-based approach were similar, but unrelated individuals have a slight decrease in R0 and a slight increase in R1 and KING-robust kinship, compared to the SFS-based approach. This makes unrelated individuals appear slightly more related than expected for unrelated individuals from a homogenous population. However, despite this bias, it is still possible to correctly classify of all the pairs to their relationship category, suggesting that the IBS-based approach can be used when not enough information is available for the SFS-based approach. The chromosome block-jackknife estimates of uncertainty for the genotype likelihood-based methods were small, and varied by relationship type, with the pair of full siblings having the most uncertainty in R0, R1, and KING-robust kinship.

We also calculated the three statistics from the high-quality phased genotypes for the same five individuals available from the 1000 Genomes Project Phase 3 (Figure 4, “1000G sites”) to see how well the two genotype-likelihood-based approaches applied to low-depth sequencing data perform compared to direct calculations from high-quality genotype data for the same samples. In this comparison, results obtained by using the genotype likelihood-based approaches applied to low-depth sequencing data are close to those obtained from the high-quality genotypes for all the pairs (Figure 4).

Finally, we also made R1-R0 plot and the R1-KING-robust kinship plots based on genotypes that we obtained through a standard genotype calling procedure from the raw read data. We did this to investigate if the genotype likelihood-based approaches are necessary or one could just as well use genotypes called from the ∼4x data. As expected, genotype calling had a large negative effect on the outcome; in the resulting R1-R0 and R1-KING-robust kinship plots the half siblings appear within the range of expected values for first cousins and both the parent-offspring and full sibling pairs appear within the range of expected values for half siblings (Figure 4, “called genotypes”). These results demonstrate the pitfalls of basing any relationship inferences, including R1-R0 and R1-KING-robust kinship plots, on genotypes called from low-depth data. Notably, this is also the case for the methods presented in Rosenberg (2006) and Manichaikul *et al*. (2010) (Figure 5A). This clearly demonstrate that, with ∼4x sequencing data, calling genotypes without external information, such as an imputation reference panel, is not a good alternative to a genotype likelihood-based approach. This implies that that software packages designed to work only on genotype data, such as KING, should not be used on data like this.

**Figure 5.**
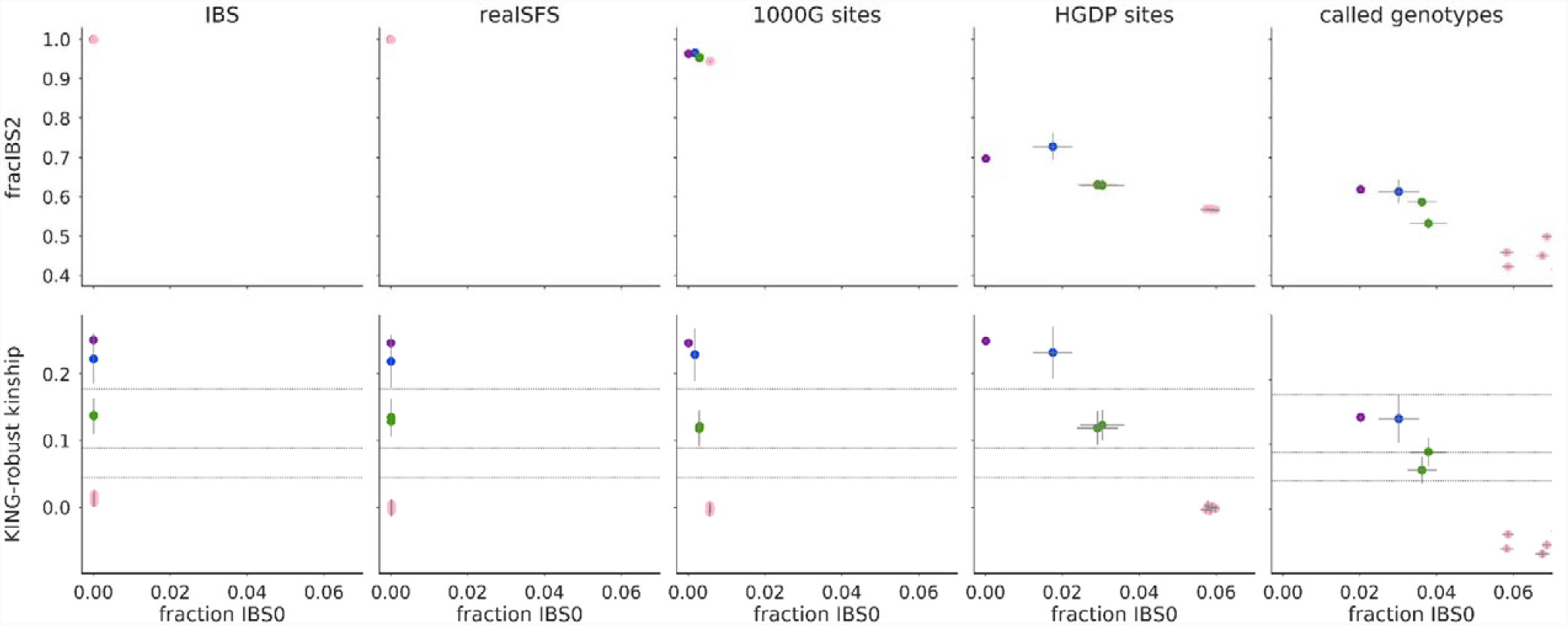
Plots of two alternate frequency-free methods for different subsets of SNP sites. **Top)** Results from applying the plotting approach from Rosenberg (2006) to pairs of the same five LWK individuals for five different analysis methods: 1) IBS: reference-free estimation from (∼4x) 1000G bam files, 2) realSFS: site-frequency spectrum based estimation from (∼4x) 1000G bam files, 3) 1000G sites: curated 1000G genotypes from the 1000G project, 4) HGDP sites: curated 1000G genotypes at sites that overlap with the Illumina 650Y array used for the HGDP, and 5) called genotypes: genotype called de novo from (∼4x) 1000G bam files. Pairs are colored by their true relationship status, as in Figure 3. Fraction IBS0/IBS2 are the overall fraction of sites that are IBS0/IBS2, respectively. Gray lines centered on each point show confidence intervals (+/-2 SE) based on a chromosome jackknife. **Bottom)** Results from applying the KING-robust based approach to the same pairs of LWK individuals using the same five different analysis methods as in A. The horizontal black lines show the kinship thresholds used to distinguish unrelated (UR), first-cousins (C1) half siblings (HS), full siblings (FS) and (PO) following (Manichaikul *et al*. 2010) from bottom to top, respectively. Like in A, thin grey lines centered on each point show confidence intervals (+/-2 SE) estimated using chromosome jackknife.

### Assessing the effect of SNP ascertainment

To assess the effect of SNP ascertainment, we applied the new method to three different subsets of data from the five 1000 Genomes individuals. The results for each of the three ascertainment schemes (all sites covered by sequencing data, 1000 Genomes release sites, Illumina 650Y SNP array sites) are similar (left four panels, Figure 4), showing SNP ascertainment does not have a large effect.

For comparison, we performed the same assessment of the methods presented in Rosenberg (2006) and Manichaikul *et al*. (2010) by constructing scatterplots of the same type as those shown in their papers (Figure 5). This revealed that both these other methods are much more affected by ascertainment than the method proposed here. In particular, the Rosenberg method is affected on both its x-axis (IBS0) and its y-axis (IBS2), which means that the expected region of the plot for each relationship will be different for different data sets (top part of Figure 5). The method presented in Manichaikul *et al*. 2010 is affected by the SNP ascertainment mainly on its x-axis (IBS0, bottom part of Figure 5). Therefore, the ascertainment mainly affects the ability to distinguish between parent-offspring and full siblings, since the y-axis, which is only slightly affected by ascertainment, is the kinship coefficient, which can be used to distinguish between most close relationships except for parent-offspring and full siblings. The x-axis, IBS0, is included in part to help make the distinction between PO and FS (Manichaikul *et al*. 2010), but this ability is clearly affected by SNP ascertainment (bottom part of Figure 5).

To further explore the effect of SNP ascertainment on the new method, we also performed analyses of the previously mentioned simulated of data from population with a constant population size. This time we only analyzed SNPs with MAF above 5% and compared the results to the results for the full dataset. This confirmed the results from the real data analyses: SNP ascertainment does change the values a bit compared to when all sites are analyzed, however the change is limited (Figure 2). This is well in line with the fact that we got very similar results for the simulated data from three populations with quite different population size histories and consequently different allele frequency spectra. Indeed, the effect of population size decline is similar to that of ascertaining for common SNPs, which makes sense because population decline is known to lead to a skew in the allele frequency spectrum towards more common SNPs.

### Assessing the effect of a limited number of sites

Genome-wide shotgun sequencing data, as is available for the 1000G individuals, is not available for all species. Studies may instead have RADseq or similar data, covering only a fraction of genomic sites. To assess to what extent the new method can be used to analyze such data sets, we performed analyses of subsets of the 1000G data, constructed to mimic RAD sequencing data. Specifically, we analyzed four subsets that consisted of the sites from 10k 50k, 100k and 250k 200bp windows, respectively. For all but the smallest data subset, the point estimates were similar to those obtained using the full dataset, showing the method is robust to reducing the number of sites with ∼4x coverage (Figure 6, supplemental file 1). This suggests that even with the reduced number of sites tested, there was sufficient data to characterize the genome-wide mean IBD fractions for both closely related and unrelated pairs. The uncertainty in the estimates, as estimated by a chromosome jackknife, increased with fewer sites, but the effect was limited, suggesting the biological variation in IBD sharing across chromosomes was larger than sampling variance across the examined sites.

**Figure 6.**
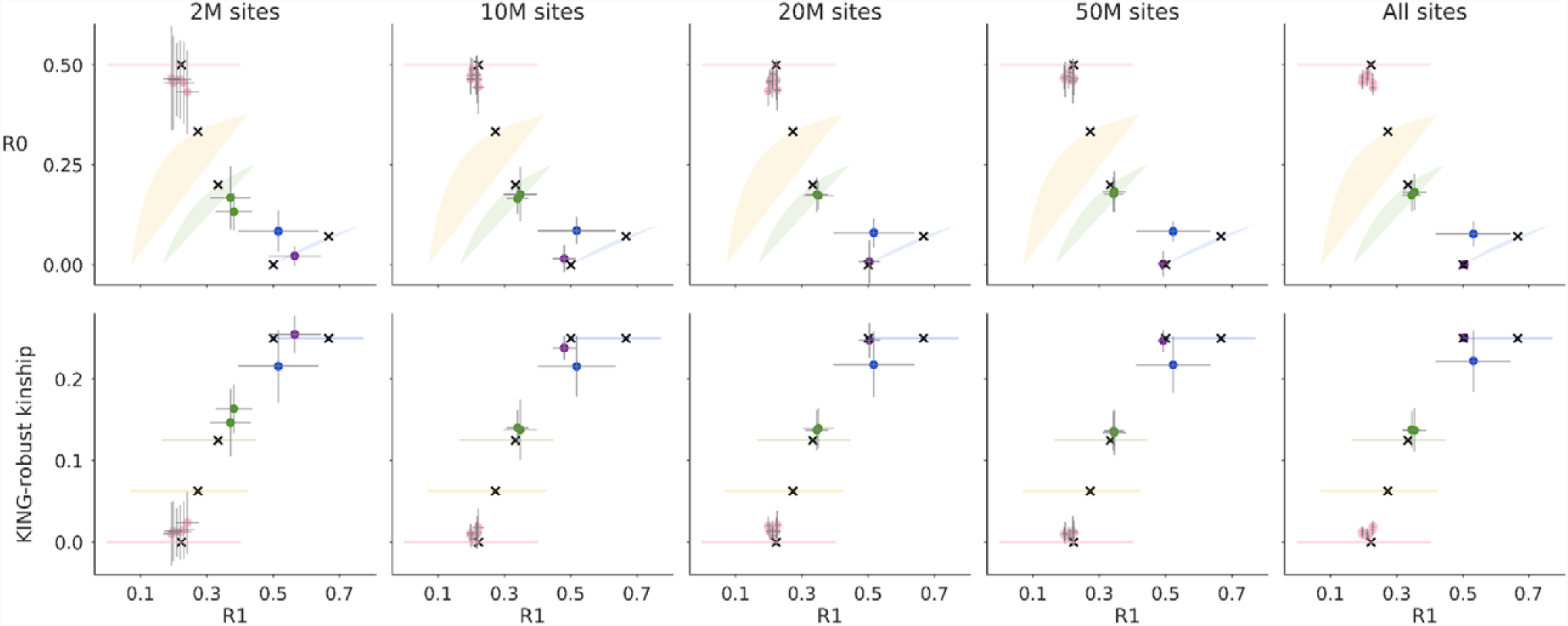
The effect on estimates of R0, R1, and KING-robust kinship from reducing the number of sites analyzed from the same five LWK individuals as in figures 4 and 5. Each point shows the point estimate and error bars show +/-2SE estimated using chromosome jackknife. Each column shows results for different numbers of examined basepairs. Pairs of individuals are colored by their true relationship. These plots show the results of the IBS-based method, see supplemental for results from the SFS-based approach. Colored shaded areas/lines show the theoretically derived range of expected values for specific relationship categories from Figure 2 and black ‘X’s show the values for a pair of individuals simulated under a neutral demography from Figure 2.

## Discussion

We have presented a simple new method for inferring if and how two individuals are related based solely on genetic data from the two target individuals. This new method ways in which it can also be applied directly to sequencing data via genotype likelihoods. And importantly, we show that by using these, the method provides useful results when applied to ∼4x sequencing data as well as RADseq like subsets of such data. All of this combined implies that – unlike previous methods - this new method can be used even if all you have is low-depth sequencing data from a few individuals from a species without a reference genome.

### Comparison to similar methods

The new method is based on plotting two statistics, R0 and KING-robust kinship against a third, new statistic R1. The R0 statistic similar to the test statistic proposed by Lee (2003) to test for relatedness. The only differences are that the numerator and denominator are flipped and that E, the proportion of sites where both individuals are heterozygous, is included in the denominator in the statistic defined by Lee but absent in R0. R0 is also similar to the pairwise population concordance (PPC) statistic in PLINK (Purcell *et al*. 2007), a test of if the genotypes of a pair of individuals have more IBS0 sites than two unrelated individuals with the same ancestry are expected to have, signaling they have different ancestries. The KING-robust kinship estimator was proposed by Manichaikul *et al*. (2010) and implemented for genotype data in the program KING. Here we extend it to estimation directly from sequencing data. Notably, our results suggest that this extension is vital for successful application to low-depth sequencing data, because estimates based on genotypes called from low-depth sequencing data are very poor (rightmost panels of Figure 4 and 5), which makes programs like KING inappropriate to apply to such data. And importantly this extension can also be used for similar statistics and thus makes existing methods based on such statistics, like KING-robust kinship, more widely applicable.

However, this extension is not the only contribution of this study. Another key new contribution is provide an alternative to the IBS0 statistic (the proportion of sites where the two individuals share zero alleles IBS) that was utilized by Rosenberg (2006) and Manichaikul *et al*. (2010). As we have shown, the fraction of sites that are IBS0 or IBS2 is very sensitive SNP ascertainment, meaning that results are only comparable within each different ascertainment scheme. The method from Rosenberg (2006), where IBS0 is combined with IBS2, is difficult, if not impossible, to use for relatedness inference because the fraction of sites that are IBS2 and IBS0 varies so wildly across different ascertainment schemes, such as between SNP arrays and sequencing data,,,. On the other hand, KING-robust kinship is still very useful, but it loses the ability to distinguish between parent-offspring and full siblings, as IBS0 was used for this. Due to this sensitivity to ascertainment, samples cannot be analyzed in isolation and must be placed in the context of other samples with known relationships and the same ascertainment scheme. This requirement makes it difficult to apply these previous methods to ancient humans or other species with limited sample sizes.

In contrast, the ability to identify relatives based on expected values is maintained in the new method, regardless of ascertainment scheme due to the use of R0, instead of IBS0, which makes the new method robust to SNP ascertainment. Parent-offspring pairs tend to have an R0 estimate extremely close to 0, making them particularly easy to identify via the R1-R0 plot. The R1-KING-robust kinship plot, on the other hand, has the appealing aspect that the kinship axis has a biological interpretation, defined as the probability that two alleles sampled at random from two individuals are identical by descent. Hence the two plots types, R1-R0 and R1-KING-robust kinship, each have their advantages. Finally, it is worth noticing that the two plots types seem to work better than a range of other plots constructed from similar ratio statistics that we explored (Supplementary Figure S6).

### Limitations and applications

While the new method provides substantial advantages over previous methods in situations with limited data, it does have some limitations. First, like most other relatedness inference methods, like PLINK, the proposed method assumes that the individuals are not inbred and that they originate from the same homogenous, population. And like many other relationship inference methods, it is not necessarily robust to violations of these assumptions. Previous studies have shown the effect of population structure and admixture on relatedness inference is complex and can potentially lead to bias in either direction depending on the circumstances, and this is true even for KING, which was developed to be robust to population structure (Thornton *et al*. 2012; Conomos *et al*. 2016; Ramstetter *et al*. 2017). Specific methods have been developed to correct for admixture when the allele frequencies in the admixing populations are known (e.g. Thornton *et al*. 2012; Moltke and Albrechtsen 2014), or enough samples are available (Conomos *et al*. 2016; Dou *et al*. 2017). But since these methods work by exploiting knowledge about allele frequencies or access to many samples for their correction; the pairwise R0, R1 and KING-robust kinship statistics cannot be corrected in a similar manner. However, we note that Lee (2003) showed that the statistic he proposed for testing for relatedness can also be used to detect if two unrelated samples are not from the same homogeneous population. If this is the case, the Lee’s statistic will be significantly smaller than 2/3; and equivalently R0 will be significantly above 0.5, which may be useful when interpreting Ro, R1, KING-kinship plots in the presence of admixture or population structure more generally. Regarding inbreeding, one potential way to assess if one of the individuals is inbred is to compare heterozygosities across individuals; non-inbred and non-admixed individuals from the same population should have similar heterozygosity, so marked heterozygosity differences can be a warning signal.

A second limitation, which is shared with other relatedness estimation methods, is that there is significant biological variation in the amount of IBD sharing between relatives with the same pedigree relationship due to randomness inherent in the process of recombination (Hill 1993; Rasmuson 1993). For humans, this means that a pair of relatives, say first cousins, will sometimes share less of their genomes IBD than another pair with a more distant pedigree relationship, say second cousins. This makes classification into specific relationships difficult. The degree of biological variation in IBD sharing between relatives varies across species and can even differ between sexes due to sex-specific recombination patterns. This makes it difficult to provide general guidance appropriate for all species. In general, species with more chromosomes and more recombination will have less variation in IBD sharing for a defined pedigree relationship, making it easier to distinguish among various potential relationship categories. To quantify this uncertainty, we propose a chromosomal bootstrap procedure that can be used if reads can be assigned to chromosomes.

Biological variation in IBD sharing is also related to the estimation and interpretation of confidence intervals on statistics like R0, R1, and KING-robust kinship. Relatedness and limited recombination also cause correlation between sites in the genome, due to shared IBD segments and LD. This correlation between sites increases the variance in the estimates of these statistics in a way that can be difficult to fully account for when computing confidence intervals. For statistics that test for introgression such as the D-statistic (Patterson *et al*. 2012), where the main concern is correlation due to LD, a block jackknife, leaving out contiguous blocks (e.g. 5Mb) is a common approach. When considering relatedness, we want to compare our estimates to the expectations of each relationship category. Since shared IBD segments can be much longer than the range of LD we propose a more appropriate chromosome-jackknife. In either case, a jackknife (or bootstrap) over single sites will fail to provide a confidence interval that accounts for the non-independence of the sites. For more discussion on this topic see Thompson (2013). Unfortunately, this means that it is difficult to provide the most appropriate confidence intervals when no information about genomic positions is available.

Despite these limitations, we believe that the results presented here suggest the new method constitutes a helpful new tool for relatedness inference for studies with limited data. Identifying related samples is a crucial step in nearly any genetic analysis and can also reveal other problems such as duplicate samples or cross contamination of genetic material. Removing the requirements to specify allele frequencies and to have accurate genotypes has the potential allow the identification of relatives even in small studies of non-model species or ancient samples. These types of studies do not currently have many good options to address relatedness.

## Data Accessibility

The IBS method is available at: http://www.popgen.dk/angsd/index.php/Genotype_Distribution

The data sets used are publicly available.

The HGDP SNP array data are available at: ftp.cephb.fr/hgdp_supp1

The 1000G phase 3 aligned sequencing data are available at: ftp://ftp.1000genomes.ebi.ac.uk/vol1/ftp/phase3/data

The 1000G phase 3 called genotypes are available at: ftp://ftp.1000genomes.ebi.ac.uk/vol1/ftp/release/20130502/

## Author Contributions

AA, IM, and RKW designed the research. RKW performed research, with input from IM and AA. IM and RKM wrote the paper, with help from AA.

