## Supplementary Materials for "Allele frequency-free inference of close familial relationships from genotypes or low depth sequencing data" (from another version of this preprint: version 2, 2018-03-20)

March 20, 2018

2.1.5 Derivation of the expected values of the KING-robust kinship estimator . . 12

### 1 Supplemental Figures

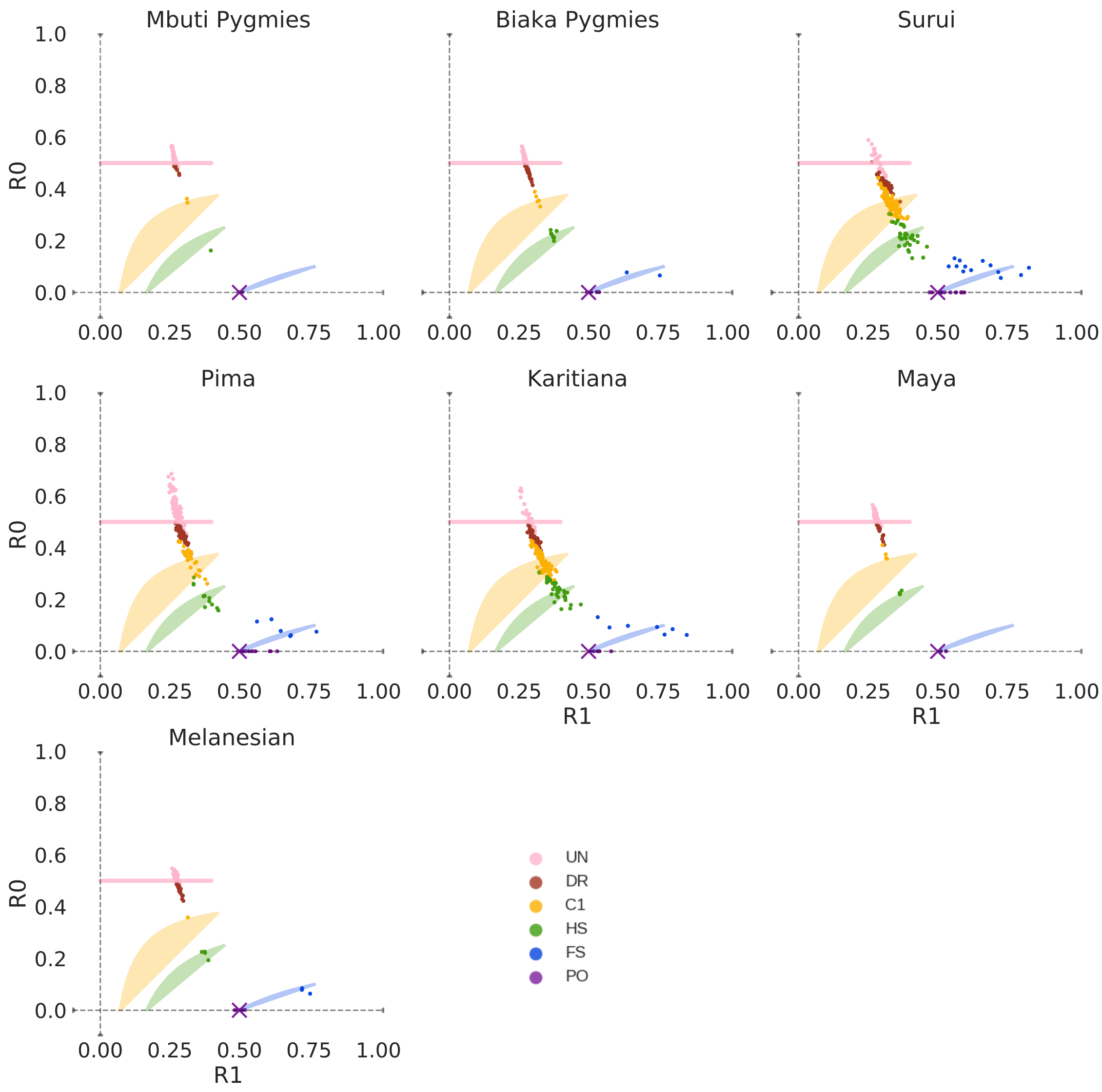

Figure S1: Scatterplot of R0 and R1 per HGDP population. Each pair of individuals within each population is represented by a point, which is colored according to the relationship category inferred using an allele frequency-based approach. Shaded areas and lines show the expected ranges for specific relationship categories, as in Figures 2 and 3 in the main text. PO = parent-offspring, FS = full sibling, HS = half sibling, C1 = first cousin, DR = distantly related, UN = unrelated.

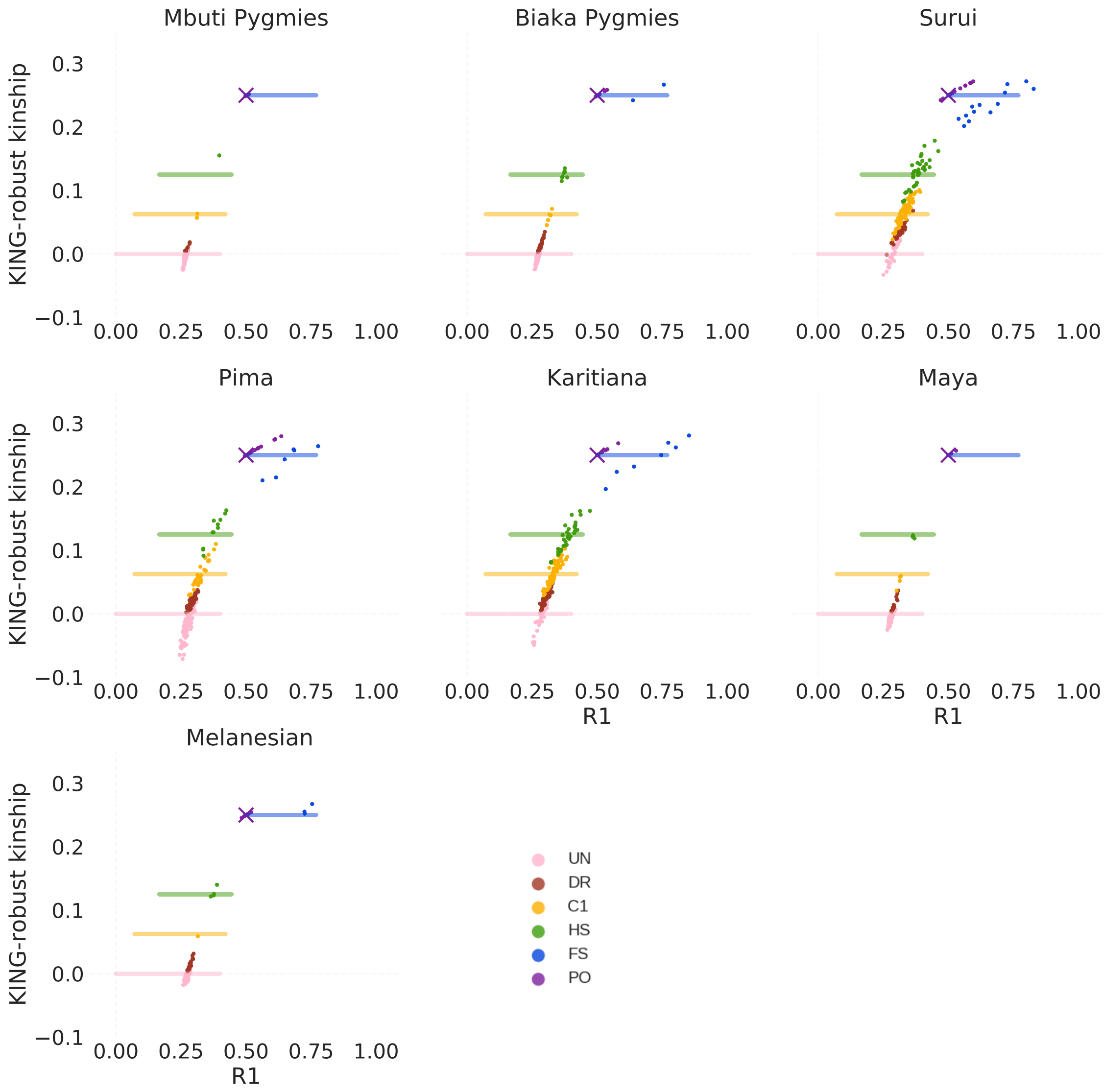

Figure S2: Scatterplot of R1 and KING-robust kinship per HGDP population. Each pair of individuals within each population is represented by a point, which is colored according to the relationship category inferred using an allele frequency-based approach. Shaded areas and lines show the expected ranges for specific relationship categories, as in Figures 2 and 3 in the main text. PO = parent-offspring, FS = full sibling, HS = half sibling, C1 = first cousin, DR = distantly related, UN = unrelated.

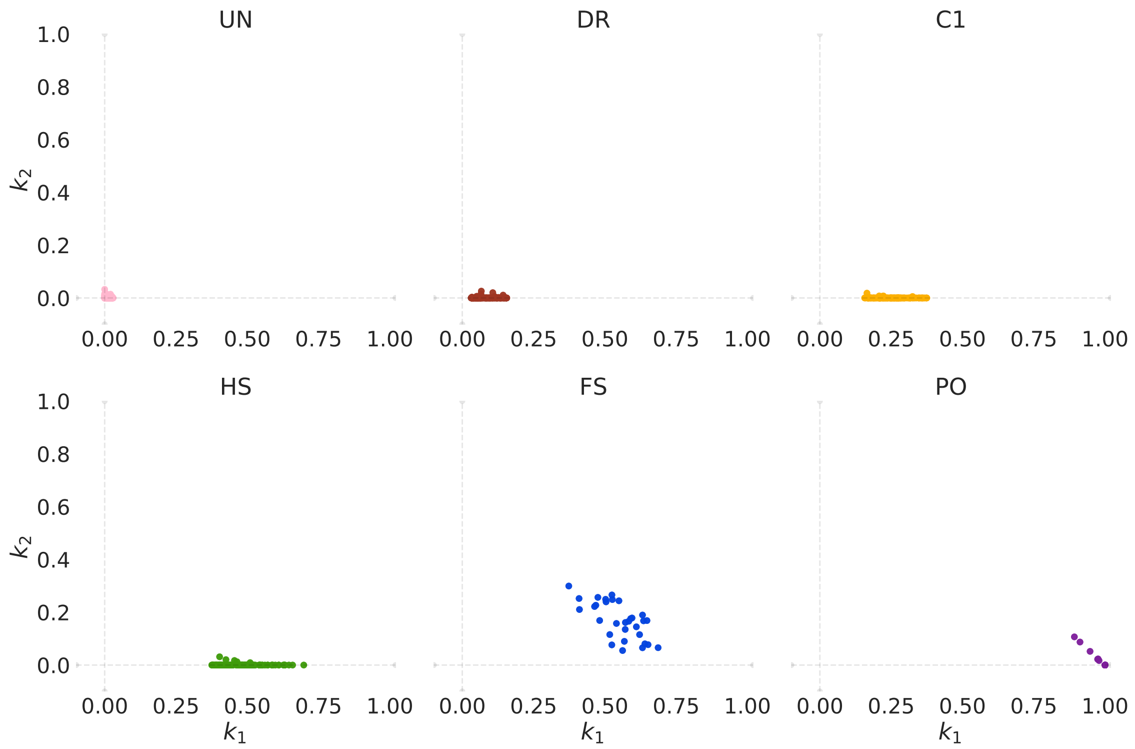

Figure S3: Scatterplot of the two relatedness coefficients $k_1$ (denoted Z1 in the output of PLINK) and $k_2$ (denoted Z2 in the output of PLINK) for each relationship category in the HGDP data. Estimates of the two relatedness coefficients are from the allele frequency-based approach implemented in PLINK. Each pair of individuals within each population is represented by a point, here they are paneled by the inferred relationship category: PO = parent-offspring, FS = full sibling, HS = half sibling, C1 = first cousin, DR = distantly related, UN = unrelated. Also see figure 3 in the main text.

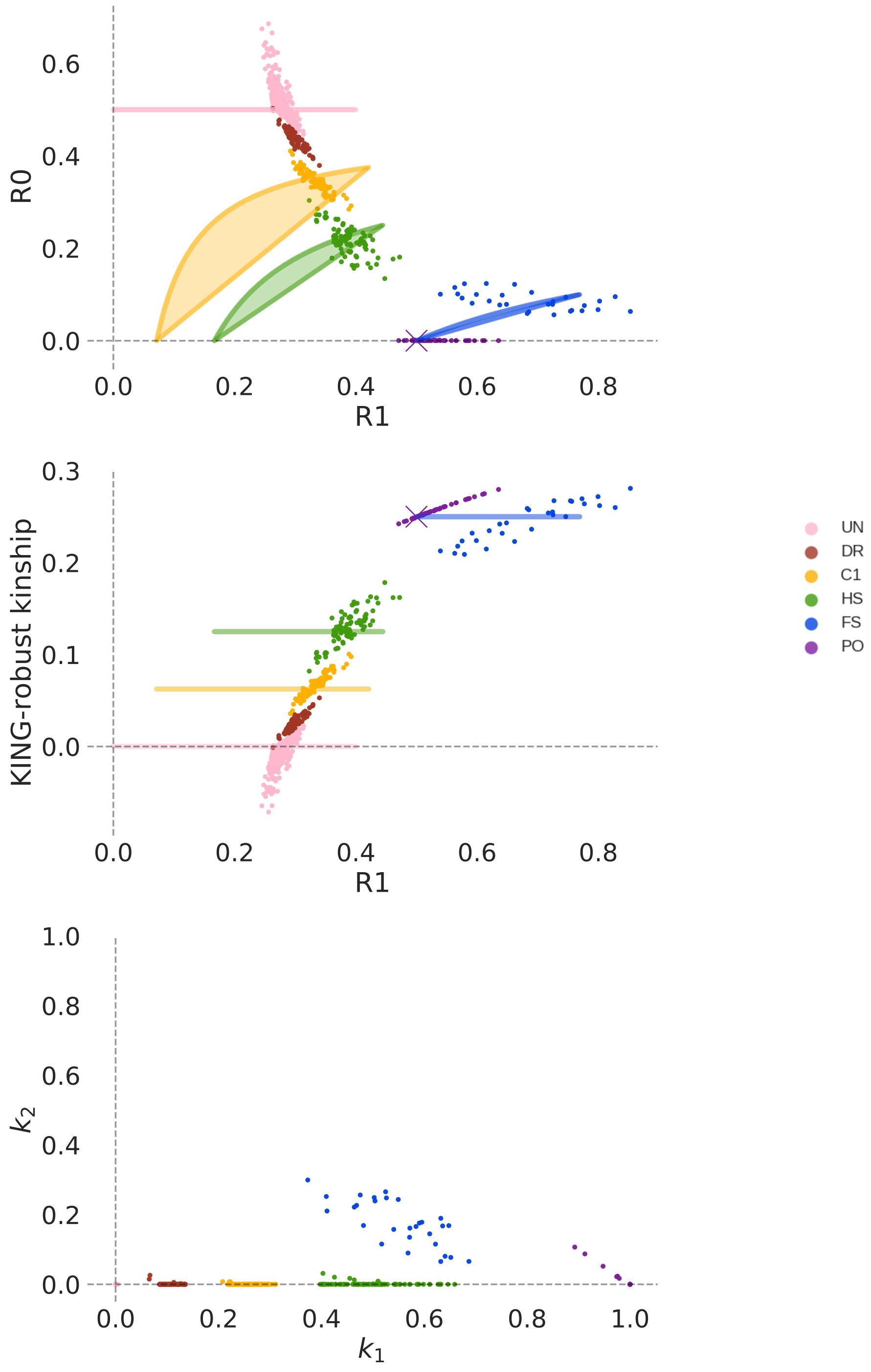

Figure S4: Pairwise scatterplots of R1, R0, and KING-robust kinship, and also $k_1$ vs $k_2$ when difficult to call relationships are excluded. Each pair of individuals within each population is represented by a point, which is colored according to the relationship category inferred using an allele frequency-based approach. Shaded areas and lines show the expected ranges for specific relationship categories, as in Figures 2 and 3 in the main text. PO = parent-offspring, FS = full sibling, HS = half sibling, C1 = first cousin, DR = distantly related, UN = unrelated. Relationships were deemed difficult to call when the PLINK kinship was within 0.02 of the cutoff between two relationship categories.

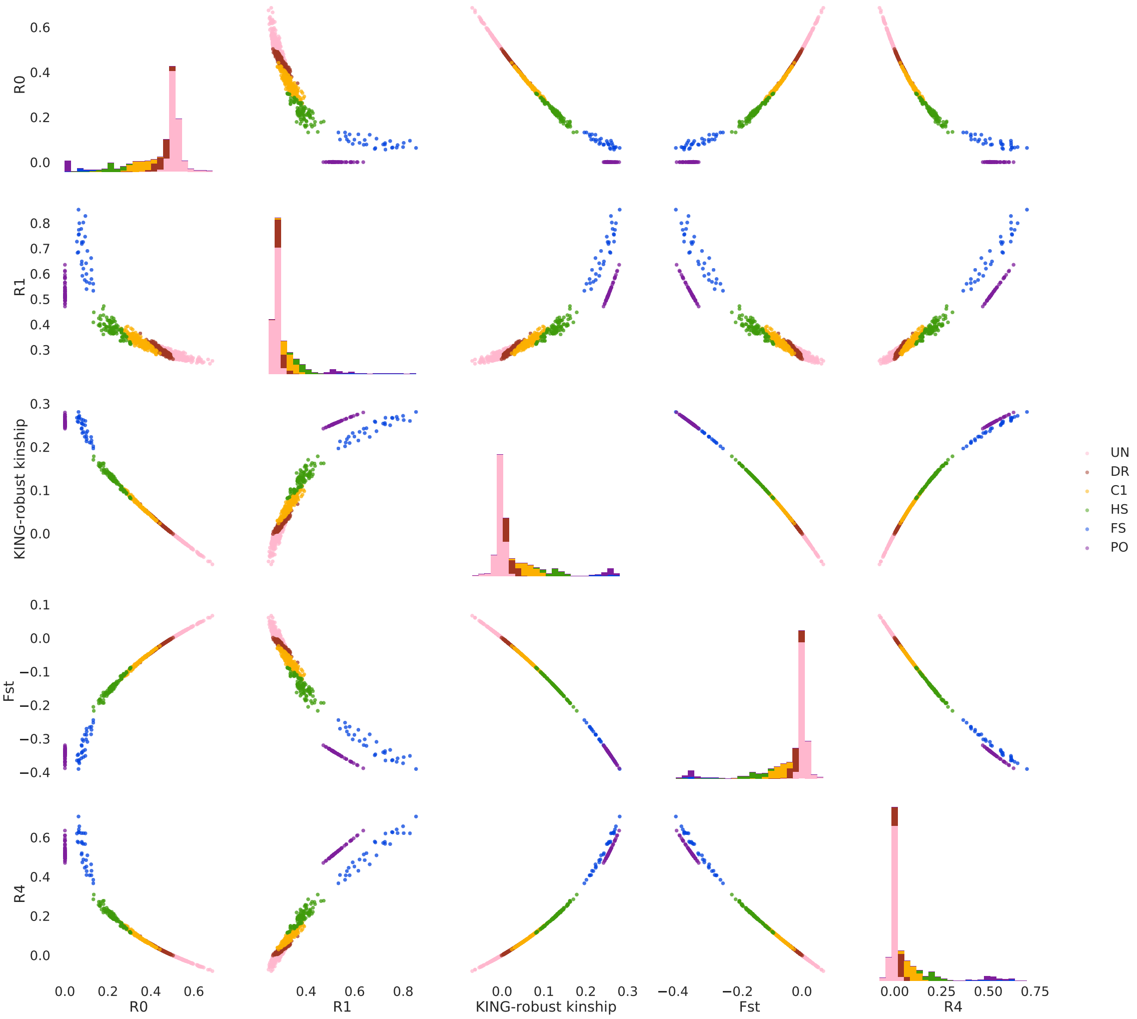

Figure S5: Scatterplots of R0, R1, KING-robust kinship, $F_{ST}$, and R4 for pairs of individuals within the selected HGDP populations. Each pair of individuals within each population is represented by a point, which is colored according to the relationship category inferred using an allele frequency-based approach. Histograms of each statistic are on the diagonal. It is evident that pairs of statistics reduce the overlap in expected ranges between relationship categories. Also presented here are two related ratios not discussed in the main text: $F_{ST} = \frac{2C+2G-E}{2C+2G+B+D+E+F+H}$ and R4 $= \frac{E-2C-2G}{2C+2G+B+D+F+H}$. PO = parent-offspring, FS = full sibling, HS = half sibling, C1 = first cousin, DR = distantly related, UN = unrelated.

### 2 Supplemental Texts

#### 2.1 Text S1: Derivations of the expectations of R0, R1, and KING-robust kinship

We will here derive expressions for R0, R1 and KING-robust kinship for a range of different pairwise familial relationships. Based on these expressions we will then determine the expected joint ranges of R0 and R1 as well as R0 and KING robust kinship shown in figure 2 in the main text.

##### 2.1.1 Assumptions and notation

In the below derivations will assume that we are analyzing data from two individuals, 1 and 2 that are not inbred and that are from the same homogeneous population. Additionally, we will assume that we have genotype data for $n$ sites from both individuals and that all sites $n$ sites are in Hardy-Weinberg equilibrium. In terms of notation, we will denote the two individuals' genotypes at given site $s$ as $g_1$ and $g_2$, with $g_1$,$g_2 \in \{0, 1, 2\}$ corresponding to the number of copies of a specified allele (e.g., the derived allele) carried by individual 1 and 2, respectively. Also, we will denote the true frequency of the specified allele at site $s$ as $f$. Furthermore, we will use the capital letters $A$ through $I$ to denote the probability of each of the nine different genotype pairs as shown in figure 1 in the main text. So e.g., $A$ denotes the probability that both individuals have the genotype 0 at a site. Finally, we will use $k_0$, $k_1$, and $k_2$ to denote the probability that the two individuals share 0, 1 or 2 alleles identical-by-descent (IBD), respectively. The expected values of $K = (k_0, k_1, k_2)$ for different familial relationship can be seen in table S1.

Table S1: Expected $K = (k_0, k_1, k_2)$ for different relationship categories.

| Relationship | $k_0$ | $k_1$ | $k_2$ |
|---|---|---|---|
| Monozygotic twins (MZ) | 0 | 0 | 1 |
| Parent-offspring (PO) | 0 | 1 | 0 |
| Full siblings (FS) | $\frac{1}{4}$ | $\frac{1}{2}$ | $\frac{1}{4}$ |
| Half siblings/avuncular/grandparent-grandchild (HS) | $\frac{1}{2}$ | $\frac{1}{2}$ | 0 |
| First cousins (C1) | $\frac{3}{4}$ | $\frac{1}{4}$ | 0 |
| Second cousins (C2) | $\frac{15}{16}$ | $\frac{1}{16}$ | 0 |
| Unrelated (UR) | 1 | 0 | 0 |

##### 2.1.2 Derivations of A through I

To derive the expected value of R0, R1 and KING-robust kinship for different familial relationship pairs, we first derive expressions for A to I. To do this we note that in general it must hold that in site $s$:

$$\begin{aligned} P(g_1, g_2 | K = (k_0, k_1, k_2), f) &= P(g_1|f)P(g_2|g_1, K = (k_0, k_1, k_2), f) \\ &= P(g_1|f) \sum_{z\in\{0,1,2\}} P(Z = z|K = (k_0, k_1, k_2))P(g_2|g_1, Z = z, f) \\ &= P(g_1|f) \sum_{z\in\{0,1,2\}} k_z P(g_2|g_1, Z = z, f) \end{aligned}$$

where $Z$ is an indicator of whether the two individuals share 0, 1 or 2 alleles IBD in a given site. Since we are assuming that the two individuals 1 and 2 are not inbred and that all sites are in Hardy-Weinberg equilibrium, the values of $P(g_1|f)$, $P(g_2|g_1, Z = 0, f)$, $P(g_2|g_1, Z = 1, f)$ and $P(g_2|g_1, Z = 2, f)$ must be those given in tables S2 to S5.

Based on this, we can derive expressions for A through I for an arbitrary degree of relatedness specified by $K$:

$$\begin{aligned}
A &= P(g_1 = 0, g_2 = 0 | K = (k_0, k_1, k_2), f) \\
&= P(g_1 = 0 | f) \sum_{z \in \{0,1,2\}} k_z P(g_2 = 0 | g_1 = 0, Z = z, f) \\
&= f^2 (k_0 f^2 + k_1 f + k_2 1) \\
&= k_0 f^4 + k_1 f^3 + k_2 f^2
\end{aligned}$$

$$\begin{aligned}
B &= P(g_1 = 1, g_2 = 0 | K = (k_0, k_1, k_2), f) \\
&= P(g_1 = 1 | f) \sum_{z \in \{0,1,2\}} k_z P(g_2 = 0 | g_1 = 1, Z = z, f) \\
&= 2f(1-f)(k_0 f^2 + k_1 \frac{f}{2} + k_2 0) \\
&= k_0 2 f^3 (1-f) + k_1 f^2 (1-f)
\end{aligned}$$

$$\begin{aligned}
C &= P(g_1 = 2, g_2 = 0 | K = (k_0, k_1, k_2), f) \\
&= P(g_1 = 2 | f) \sum_{z \in \{0,1,2\}} k_z P(g_2 = 0 | g_1 = 2, Z = z, f) \\
&= (1-f)^2 (k_0 f^2 + k_1 0 + k_2 0) \\
&= k_0 f^2 (1-f)^2
\end{aligned}$$

$$\begin{aligned}
D &= P(g_1 = 0, g_2 = 1 | K = (k_0, k_1, k_2), f) \\
&= P(g_1 = 0 | f) \sum_{z \in \{0,1,2\}} k_z P(g_2 = 1 | g_1 = 0, Z = z, f) \\
&= f^2 (k_0 2 f (1-f) + k_1 (1-f) + k_2 0) \\
&= k_0 2 f^3 (1-f) + k_1 f^2 (1-f) \\
&= B
\end{aligned}$$

$$\begin{aligned}
E &= P(g_1 = 1, g_2 = 1 | K = (k_0, k_1, k_2), f) \\
&= P(g_1 = 1 | f) \sum_{z \in \{0,1,2\}} k_z P(g_2 = 1 | g_1 = 1, Z = z, f) \\
&= 2f(1-f)(k_0 2 f (1-f) + k_1 \frac{1}{2} + k_2 1) \\
&= k_0 4 f^2 (1-f)^2 + k_1 f (1-f) + k_2 2 f (1-f)
\end{aligned}$$

$$\begin{aligned}
F &= P(g_1 = 2, g_2 = 1 | K = (k_0, k_1, k_2), f) \\
&= P(g_1 = 2 | f) \sum_{z \in \{0,1,2\}} k_z P(g_2 = 1 | g_1 = 2, Z = z, f) \\
&= (1-f)^2 (k_0 2 f (1-f) + k_1 f + k_2 0) \\
&= k_0 2 f (1-f)^3 + k_1 f (1-f)^2
\end{aligned}$$

$$
\begin{aligned}
G &= P(g_1 = 0, g_2 = 2|K = (k_0, k_1, k_2), f) \\
&= P(g_1 = 0|f) \sum_{z\in\{0,1,2\}} k_z P(g_2 = 2|g_1 = 0, Z = z, f) \\
&= f^2(k_0(1-f)^2 + k_1 0 + k_2 0) \\
&= k_0 f^2(1-f)^2 \\
&= C
\end{aligned}
$$

$$
\begin{aligned}
H &= P(g_1 = 1, g_2 = 2|K = (k_0, k_1, k_2), f) \\
&= P(g_1 = 1|f) \sum_{z\in\{0,1,2\}} k_z P(g_2 = 2|g_1 = 1, Z = z, f) \\
&= 2f(1-f)(k_0(1-f)^2 + k_1\frac{(1-f)}{2} + k_2 0) \\
&= k_0 2f(1-f)^3 + k_1 f(1-f)^2 \\
&= F
\end{aligned}
$$

$$
\begin{aligned}
I &= P(g_1 = 2, g_2 = 2|K = (k_0, k_1, k_2), f) \\
&= P(g_1 = 2|f) \sum_{z\in\{0,1,2\}} k_z P(g_2 = 2|g_1 = 2, Z = z, f) \\
&= (1-f)^2(k_0(1-f)^2 + k_1(1-f) + k_2 1) \\
&= k_0(1-f)^4 + k_1(1-f)^3 + k_2(1-f)^2
\end{aligned}
$$

Table S2: $P(g_1|f)$.

| $P(g_1 = 0|f)$ | $P(g_1 = 1|f)$ | $P(g_1 = 2|f)$ |
|---|---|---|
| $f^2$ | $2f(1-f)$ | $(1-f)^2$ |

Table S3: $P(g_2|g_1, Z = 0, f)$.

| | $g_2 = 0$ | $g_2 = 1$ | $g_2 = 2$ |
|---|---|---|---|
| $g_1 = 0$ | $f^2$ | $2f(1-f)$ | $(1-f)^2$ |
| $g_1 = 1$ | $f^2$ | $2f(1-f)$ | $(1-f)^2$ |
| $g_1 = 2$ | $f^2$ | $2f(1-f)$ | $(1-f)^2$ |

Table S4: $P(g_2|g_1, Z = 1, f)$.

| | $g_2 = 0$ | $g_2 = 1$ | $g_2 = 2$ |
|---|---|---|---|
| $g_1 = 0$ | $f$ | $(1-f)$ | $0$ |
| $g_1 = 1$ | $\frac{f}{2}$ | $\frac{1}{2}$ | $\frac{1-f}{2}$ |
| $g_1 = 2$ | $0$ | $f$ | $1-f$ |

Table S5: $P(g_2|g_1, Z = 2, f)$.

| | $g_2 = 0$ | $g_2 = 1$ | $g_2 = 2$ |
|---|---|---|---|
| $g_1 = 0$ | 1 | 0 | 0 |
| $g_1 = 1$ | 0 | 1 | 0 |
| $g_1 = 2$ | 0 | 0 | 1 |

##### 2.1.3 Derivation of the expected values of R0

With the above expressions for A through I, we can derive an expectation of R0 for different relationships. We do this by first noting that:

$$\begin{aligned} R0 &= \frac{C+G}{E} \\ &= \frac{2C}{E} \\ &= \frac{2(k_0 f^2(1-f)^2)}{k_0 4f^2(1-f)^2 + k_1 f(1-f) + k_2 2f(1-f)} \\ &= \frac{2k_0 f(1-f)}{k_0 4f(1-f) + k_1 + 2k_2} \end{aligned}$$

Inserting the expected values of $k_0, k_1$ and $k_2$ from table S1 into this formula we get that if individuals 1 and 2 are a PO pair $R0$ is expected to be

$$\begin{aligned} R0_{PO} &= \frac{2 \times 0 f(1-f)}{0 \times 4f(1-f) + 1 + 2 \times 0} \\ &= \frac{0}{1} \\ &= 0 \end{aligned}$$

Similarly, for monozygotic twins (MZ), full siblings (FS), half siblings/avuncular/grandparent-grandchild (HS), first cousins (C1), second cousins (C2) and unrelated (UR) we expect $R0$ to be:

$$\begin{aligned} R0_{MZ} &= \frac{2 \times 0 f(1-f)}{0 \times 4f(1-f) + 0 + 2 \times 1} \\ &= 0 \end{aligned}$$

$$\begin{aligned} R0_{FS} &= \frac{2 \times \frac{1}{4} f(1-f)}{\frac{1}{4} \times 4f(1-f) + \frac{1}{2} + 2 \times \frac{1}{4}} \\ &= \frac{\frac{1}{2} f(1-f)}{f(1-f) + 1} \\ &= \frac{f(1-f)}{2f(1-f) + 2} \end{aligned}$$

which for $f \in ]0, 1[$ has the range $]0, \frac{1}{10}]$.

$$\begin{aligned} R0_{HS} &= \frac{2 \times \frac{1}{2} f(1-f)}{\frac{1}{2} \times 4f(1-f) + \frac{1}{2} + 2 \times 0} \\ &= \frac{f(1-f)}{2f(1-f) + \frac{1}{2}} \end{aligned}$$

which for $f \in ]0,1[$ has the range $]0, \frac{1}{4}]$.

$$
\begin{aligned}
R0_{C1} &= \frac{2 \times \frac{3}{4} f(1-f)}{\frac{3}{4} \times 4f(1-f) + \frac{1}{4} + 2 \times 0} \\
&= \frac{\frac{3}{2} f(1-f)}{3f(1-f) + \frac{1}{4}}
\end{aligned}
$$

which for $f \in ]0,1[$ has the range $]0, \frac{3}{8}]$.

$$
\begin{aligned}
R0_{C2} &= \frac{2 \times \frac{15}{16} \times f(1-f)}{\frac{15}{16} \times 4f(1-f) + \frac{1}{16} + 2 \times 0} \\
&= \frac{30f(1-f)}{60f(1-f) + 1}
\end{aligned}
$$

which for $f \in ]0,1[$ has the range $]0, \frac{15}{32}]$.

$$
\begin{aligned}
R0_{UR} &= \frac{2 \times 1 \times f(1-f)}{1 \times 4f(1-f) + 0 + 2 \times 0} \\
&= \frac{2f(1-f)}{4f(1-f)} \\
&= \frac{1}{2}
\end{aligned}
$$

which is constant independent of the value for $f \in ]0,1[$.

##### 2.1.4 Derivation of the expected values of R1

With the above expressions for A through I we also get that

$$
\begin{aligned}
R1 &= \frac{E}{B + C + D + F + G + H} \\
&= \frac{E}{2B + 2C + 2F} \\
&= \frac{k_0 4f^2(1-f)^2 + k_1 f(1-f) + k_2 2f(1-f)}{2(k_0 2f^3(1-f) + k_1 f^2(1-f)) + 2(k_0 f^2(1-f)^2) + 2(k_0 2f(1-f)^3 + k_1 f(1-f)^2)} \\
&= \frac{k_0 4f^2(1-f)^2 + k_1 f(1-f) + k_2 2f(1-f)}{k_0(4f^3(1-f) + 2f^2(1-f)^2 + 4f(1-f)^3) + k_1(2f^2(1-f) + 2f(1-f)^2)} \\
&= \frac{k_0 4f(1-f) + k_1 + k_2 2}{k_0(4f^2 + 2f(1-f) + 4(1-f)^2) + k_1(2f + 2(1-f))} \\
&= \frac{k_0 4f(1-f) + k_1 + k_2 2}{k_0(4 - 6f(1-f)) + k_1 2}
\end{aligned}
$$

Inserting the expected values of $k_0, k_1$ and $k_2$ from table S1 into this formula we get that if individuals $i1$ and $i2$ are a PO pair $R1$ is expected to be

$$
\begin{aligned}
R1_{PO} &= \frac{0 \times 4f(1-f) + 1 + 0 \times 2}{0 \times (4 - 6f(1-f)) + 1 \times 2} \\
&= \frac{1}{2}
\end{aligned}
$$

Similarly, for MZ, FS, HS, C1, C2 and UR we expects $R1$ to be:

$$
\begin{aligned}
R1_{MZ} &= \frac{0 \times 4f(1-f) + 0 + 1 \times 2}{0 \times (4 - 6f(1-f)) + 0 \times 2} \\
&= \infty
\end{aligned}
$$

$$
\begin{aligned}
R1_{FS} &= \frac{\frac{1}{4} \times 4f(1-f) + \frac{1}{2} + \frac{1}{4} \times 2}{\frac{1}{4} \times (4 - 6f(1-f)) + \frac{1}{2} \times 2} \\
&= \frac{f(1-f) + 1}{2 - \frac{3}{2}f(1-f)}
\end{aligned}
$$

which for $f \in ]0,1[$ has the range $]\frac{1}{2}, \frac{10}{13}]$.

$$
\begin{aligned}
R1_{HS} &= \frac{\frac{1}{2} \times 4f(1-f) + \frac{1}{2} + 0 \times 2}{\frac{1}{2} \times (4 - 6f(1-f)) + \frac{1}{2} \times 2} \\
&= \frac{2f(1-f) + \frac{1}{2}}{3 - 3f(1-f)}
\end{aligned}
$$

which for $f \in ]0,1[$ has the range $]\frac{1}{6}, \frac{4}{9}]$.

$$
\begin{aligned}
R1_{C1} &= \frac{\frac{3}{4} \times 4f(1-f) + \frac{1}{4} + 0 \times 2}{\frac{3}{4} \times (4 - 6f(1-f)) + \frac{1}{4} \times 2} \\
&= \frac{3f(1-f) + \frac{1}{4}}{\frac{7}{2} - \frac{9}{2}f(1-f)}
\end{aligned}
$$

which for $f \in ]0,1[$ has the range $]\frac{1}{14}, \frac{8}{19}]$.

$$
\begin{aligned}
R1_{C2} &= \frac{\frac{15}{16} \times 4f(1-f) + \frac{1}{16} + 0 \times 2}{\frac{15}{16} \times (4 - 6f(1-f)) + \frac{1}{16} \times 2} \\
&= \frac{\frac{60}{16}f(1-f) + \frac{1}{16}}{\frac{62}{16} - \frac{90}{16}f(1-f)}
\end{aligned}
$$

which for $f \in ]0,1[$ has the range $]\frac{1}{62}, \frac{32}{79}]$

$$
\begin{aligned}
R1_{UR} &= \frac{1 \times 4f(1-f) + 0 + 0 \times 2}{1 \times (4 - 6f(1-f)) + 0 \times 2} \\
&= \frac{4f(1-f)}{4 - 6f(1-f)} \\
&= \frac{2f(1-f)}{2 - 3f(1-f)}
\end{aligned}
$$

which for $f \in ]0,1[$ ranges from $]0, \frac{2}{5}]$.

##### 2.1.5 Derivation of the expected values of the KING-robust kinship estimator

Using the above expressions for A through I, the KING-robust kinship estimator (Manichaikul et al., 2010) can be re-written as:

$$\begin{aligned}
KING\text{-}robust\ kinship &= \frac{E-2(C+G)}{B+D+H+F+2E} \\
&= \frac{E-4C}{2(B+F+E)} \\
&= \frac{k_1 f(1-f)+k_2 2f(1-f)}{2(k_0 2f(1-f)+k_1 2f(1-f)+k_2 2f(1-f))} \\
&= \frac{k_1 f(1-f)+k_2 2f(1-f)}{4f(1-f)(k_0+k_1+k_2)} \\
&= \frac{k_1 f(1-f)+k_2 2f(1-f)}{4f(1-f)} \\
&= \frac{k_1}{4}+\frac{k_2}{2}
\end{aligned}$$

Hence, as expected, the expectation of the KING-robust kinship estimator is $\frac{k_1}{4}+\frac{k_2}{2}$ (which is the definition of kinship) regardless of the allele frequencies. Thus using the values in table S1 this means that the expected KING-robust kinship estimate is $\frac{1}{2}$ for MZ, $\frac{1}{4}$ for both PO pairs and full siblings, $\frac{1}{8}$ for HS, $\frac{1}{16}$ for C1, $\frac{1}{64}$ for C2 and 0 for unrelated pairs.

###### 2.1.6 Joint ranges of R1 and R0

Above we derived the ranges of the expectation of each of R1 and R0 for different relationships. To get the joint ranges of the two, we note that the two ratios are not independent, because E is a part of both ratios. More specifically, (R1,R0) as a function of $f \in ]0,1[$ for each of the different relationships considered here is shown by the solid lines in figure 2A in the main text. As this figure reveals, these are either single points (for PO) or concave, which means that for a combination of frequencies - and thus when more sites than one is considered - the ranges will be in the colored ranges inside the solid lines. It is important to note that these are simply ranges of expectations, because they are based on expected values of $k_0, k_1$ and $k_2$ for the different relationship. Hence, the realized values for a given pair will not necessarily lie inside the ranges shown as the realized values of $k_0, k_1$ and $k_2$ may differ from the expected values because the realized values for any related pair – except for parent off-spring and monozygotic twins – will vary around the expected values of $k_0$, $k_1$, and $k_2$ due to the randomness in the recombination process. E.g. a pair of half siblings are expected to have ($k_0$, $k_1$, $k_2$)=(0.5, 0.5, 0) but can in practice end up with e.g. ($k_0$, $k_1$, $k_2$)=(0.55, 0.45, 0) or ($k_0$, $k_1$, $k_2$)=(0.45, 0.55, 0). This will lead to values outside the expected range. In other words, the expectations derived here are expected values for our statistics in the same way as the values in table S1 are the expected values for K=($k_0$, $k_1$, $k_2$).

###### 2.1.7 Joint ranges of R1 and the KING-robust kinship estimator

Above we also derived the ranges of the expectation of each R1 and KING-robust for different relationships. We get the joint ranges from figure 2B in the main text by simply combining these. Again, it is important to note that these are simply ranges of expectations, because they are based on expected values of $k_0, k_1$ and $k_2$ for the different relationship. Hence, the realized values for a given pair will not necessarily lie inside the ranges shown as the realized values of $k_0$, $k_1$ and $k_2$ may differ from the expected values.

#### 2.2 Text S2: The IBS method

In this section, we will describe technical details of the IBS method introduced in the main text. Specifically, this method aims to infer the frequency of different genotype combinations, $p$, for a pair of individuals, 1 and 2 from directly from sequencing read data. This is done using genotype likelihoods, i.e. probabilities of the read data given different genotypes, for each of the two individuals. These genotype likelihoods better reflect the uncertainty of the true genotypes that is inherent to low depth sequencing data. Briefly, this is accomplished by summing over all possible genotypes of the two individuals and weighting the probabilities using the corresponding genotype likelihoods. To fully describe the IBS method, we introduce the following notation:

- $S$ denotes the number of sites
- $X_i = (X_i^1, X_i^2, ..., X_i^S)$ denotes the sequencing read data for individual $i$ at the $S$ sites
- $\mathcal{G} = \{AA, AC, AG, AT, CC, CG, CT, GG, GT, TT\}$ denotes the set of possible genotypes
- $Q_i^s$ denotes the (unknown) genotype of individual $i$ at site $s$ with $Q_i^s \in \mathcal{G}$
- $P(X_i^s|Q_i^s = q_i)$ is the likelihood of the genotype $q_i$ for individual $i$ at site $s$
- $p$ is the vector of the frequencies of the genotype combinations that we aim to estimate

With this notation it must hold that:

$$
\begin{aligned}
P(X_1, X_2|p) &= \prod_{s=1}^{S} P(X_1^s, X_2^s|p) \\
&= \prod_{s=1}^{S} \sum_{(q_1,q_2)\in\mathcal{G}\times\mathcal{G}} P(X_1^s, X_2^s|(Q_1^s, Q_2^s) = (q_1, q_2)) \times P((Q_1^s, Q_2^s) = (q_1, q_2)|p) \\
&= \prod_{s=1}^{S} \sum_{(q_1,q_2)\in\mathcal{G}\times\mathcal{G}} P(X_1^s|Q_1^s = q_1) \times P(X_2^s|Q_2^s = q_2) \times P((Q_1^s, Q_2^s) = (q_1, q_2)|p)
\end{aligned}
$$

where the genotype likelihoods for the two individuals can be calculated using tools like ANGSD (Korneliussen et al., 2014) and where the probability of each possible genotype combination, $P((Q_1^s, Q_2^s) = (q_1, q_2)|p)$, is simply the element of $p$ that corresponds to this genotype combination. This equation provides us with a likelihood function for the parameter, $p$, and we use this likelihood function to perform maximum likelihood estimation of $p$. In practice, this is done using an EM-algorithm which we have added to the software tool ANGSD with the name IBS.

We note that we have also added other similar models to ANGSD. The only difference between these and the model presented here is that fewer parameters are estimated, i.e. $p$ is a shorter vector, and that $P((Q_1^s, Q_2^s) = (q_1, q_2)|p)$ is defined differently as a consequence. Those other models are not used in this paper.

#### 2.3 Text S3: Example command lines

```
# make consensus - needed to make saf files
{ANGSD} -b ./data/1000G_aln/NA19042.mapped.ILLUMINA.bwa.LWK.low_coverage.20130415.
    list \
-r {CHR} -minMapQ 30 -minQ 20 -setMinDepth 3 -doFasta 2 -doCounts 1 -out ./data/
    consensus.NA19042.chr{CHR}

# make *.saf files (per individual)
{ANGSD} -b ./data/1000G_aln/NA19027.mapped.ILLUMINA.bwa.LWK.low_coverage.20130415.
    list \
-r {CHR} \
-ref ./data/1000G_aln/hs37d5.fa \
-anc ./data/consensus.NA19042.chr{CHR}.fa.gz  \
-sites ./data/1000G_aln/GEM_mappability1_75mer.angsd \
-minMapQ 30 -minQ 20 -GL 2 \
-doSaf 1 -doDepth 1 -doCounts 1 \
-out ./data/1000G_aln/saf/chromosomes/NA19027_chr{CHR}

# realSFS for each pair of individuals
{realSFS} ./data/1000G_aln/saf/chromosomes/NA19042_chr{CHR}.saf.idx ./data/1000
    G_aln/saf/chromosomes/NA19027_chr{CHR}.saf.idx -r {CHR} -P 2 -tole 1e-10 > ./
    data/1000G_aln/saf/chromosomes/NA19042_NA19027_chr{CHR}.2dsfs

# make genotype likelihood file
{ANGSD} -b ./data/1000G_aln/bamlist.all.txt \
-r {CHR} \
-sites ./data/1000G_aln/GEM_mappability1_75mer.angsd \
-minMapQ 30 -minQ 20 -GL 2 \
-doGlf 1 \
-out ./data/1000G_aln/GLF/chromosomes/chr{CHR}

# IBS
{IBS} -glf ./data/1000G_aln/GLF/chromosomes/chr{CHR}.glf.gz \
-seed {CHR} -maxSites 300000000 -model 0 \
-nInd 5 -allpairs 1 \
-outFileName ./data/1000G_aln/GLF/chromosomes/chr{CHR}.model0
```
